# Tunable, proteolytic dosage control of CRISPR-Cas systems enables precise gene therapy for dosage-sensitive disorders

**DOI:** 10.1101/2024.10.09.617463

**Authors:** Noa Katz, Connie An, Minza Haque, Sophia Mahiout, Yu-Ju Lee, Josh Tycko, Meng Zhang, Jeewoo Kang, Lacramioara Bintu, Michael C Bassik, Wei-Hsiang Huang, Xiaojing J Gao

## Abstract

The ability to modulate gene expression using modular, universal genetic tools such as CRISPR-Cas shows great potential for gene therapy. Yet the inherent stochasticity of delivery methods leads to variation in target gene expression, limiting applications in systems sensitive to dosage. Thus, we implement an incoherent feedforward loop (IFFL) based on proteolytic cleavage of dCas to reduce gene expression variability against the variability of vector delivery. We demonstrate dosage control of gene activation and repression, post-delivery tuning, and mRNA delivery compatibility using a genome-integrated marker. As a therapeutically relevant example, we then turn to the dosage-sensitive gene *RAI1,* the haploinsufficiency and triplosensitivity of which cause two genetic neurodevelopmental disorders. We demonstrate more consistent *Rai1* activation using our IFFL in wild-type mouse brains compared to conventional CRISPRa. Lastly, we deliver our IFFL to an *Rai1^+/-^* mouse model, improving the survivability of neonatal pan-brain CRISPRa-based gene therapy and rescuing multiple phenotypes.

## Introduction

For basic research and potential therapies, it is now routine to deliver genetic material into cells to change the expression level of target genes through direct expression of an exogenous gene or via tools that regulate intrinsic gene expression. To deliver such tools, vectors such as lentiviruses, adeno-associated viruses (AAVs), and lipid nanoparticles encapsulating messenger RNA (mRNA) are widely used and have emerged as leading therapeutic gene-delivery modalities.^1^ However, cargo expression stemming from such delivery methods is inherently variable, leading to differences in gene expression levels between individual cells and across animals/patients.

Maintaining target gene expression at a predefined level in the face of delivery variability would greatly improve our ability to quantitatively study genotype-phenotype relationships and enable large swaths of biomedical applications sensitive to gene dosage. For example, unnaturally high gene expression can cause unexpected phenotypes and toxicities, compromising the safety and efficacy of potential gene therapies.^2–4^ This concern is even more pertinent to dosage-sensitive genetic disorders, where both increased and decreased gene expression are pathological.^5^ Rett syndrome/*MECP2* duplication syndrome which involves the dosage-sensitive gene *MECP2* is the most well-known example.^6^ However, this phenomenon is also present in many autism spectrum and neurodevelopmental disorders such as Smith-Magenis syndrome/Potocki-Lupski Syndrome, which we will elaborate on through the rest of this paper.^7–9^ Dosage sensitivity complicates the development of potential *in vivo* gene therapies aimed at correcting such under- or overexpression. This is because, despite control over the total viral titer administered, the vector copy number each cell receives varies due to stochastic uptake and position in the tissue. High dosages carry the risk of overcorrection, but simply decreasing the total vector amount leaves more target cells untreated. Thus, we seek to bridge this critical gap between the variability of gene delivery and the need to precisely control gene expression (**Fig. 1A**).

**Fig 1.**
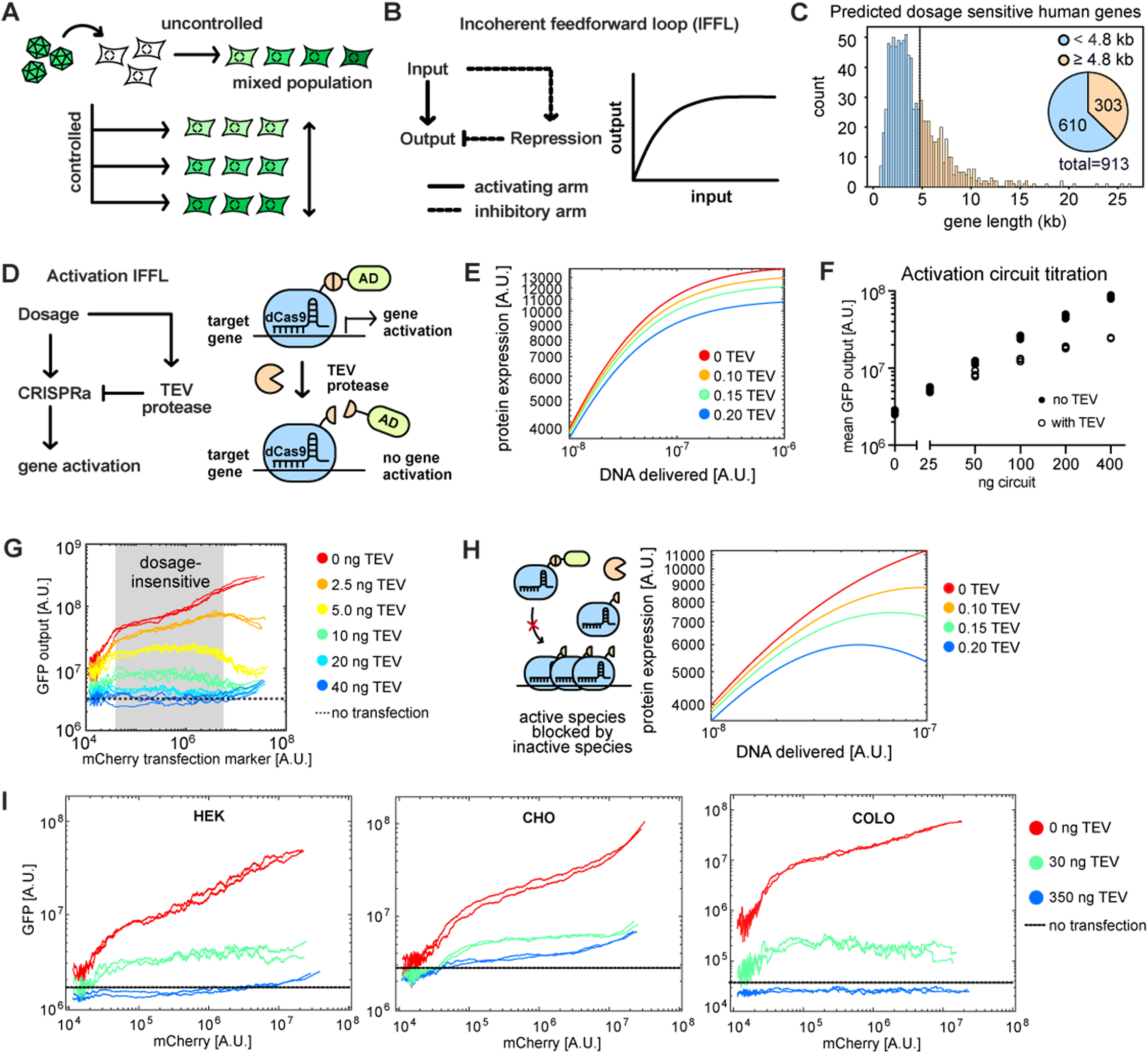
Establishing proteolytic IFFLs for gene activation using synthetic reporters. **A** Schematic of variable gene expression during gene delivery (uncontrolled) and an ideal gene delivery system which eliminates heterogeneity and tunes gene expression level (controlled). **B** Schematic of an incoherent feedforward loop. The input drives two opposing arms which allows the output to remain invariant to the input strength. **C** Histogram of sizes of human genes predicted to be dosage sensitive with a pie chart totalling the number of genes less than/greater than or equal to 4.8kb (the approximate packaging limit of an AAV). **D** Schematic of engineered activation circuit. TEV protease cleaves off the activation domain (AD) linked to dCas9, creating an IFFL with dosage as the input and CRISPRa activity as the output. **E** Simulation of the activation IFFL using ordinary differential equations. TEV fractions represent relative expression. **F** Dot plot of average GFP fluorescence (output) as a function of circuit transfected in GFP-integrated HEK293 cells. **G** Running averages of GFP fluorescence (output) as a function of mCherry fluorescence (transfection marker) with different co-transfected TEV plasmid amounts. **H** Schematic of competitive binding of active and inactive CRISPRa species and simulation of activation IFFL behavior with competitive inhibition. **I** Running average of GFP fluorescence (output) as a function of mCherry fluorescence (transfection marker) upon transient transfection of activation IFFL across three GFP-integrated cell lines. All experiments shown are based on three independent replicates.

The incoherent feedforward loop (IFFL) is a circuit motif broadly found in natural biological pathways that provides exactly this form of control.^10,11^ In an IFFL, one common input passes through an activating arm that increases the level/activity of a target gene and a parallel inhibitory arm that decreases it (**Fig. 1B**). These antagonistic arms maintain the output level against input variability: variation in the input will proportionately drive both antagonistic arms, compensating for changes and leading to a consistent output level. In the context of gene delivery, dosage or transgene copy number acts as the input and transgene expression is the output. Ideally, an engineered IFFL would (1) depend minimally on the host cell and function robustly across cell types, (2) be compatible with various delivery methods, (3) control the activation and inhibition of diverse target genes, and (4) allow for tuning of transgene expression level pre- and post-delivery. Previously engineered IFFLs directly or indirectly target an exogenously delivered gene using various inhibition factors such as transcriptional repressors, antagonists, and RNAi.^12–20^ While all have achieved impressive dosage control, the most prevalent mechanism of repression is RNAi. However, this strategy has variable IFFL performance across cell types and can saturate the endogenous RNAi machinery^14,16,18^, is not compatible with mRNA vectors (a promising emerging modality given its superior safety than more conventional DNA vectors), and has not yet demonstrated post-delivery tuning or gene repression control.

To meet these criteria, we propose a novel proteolytic cleavage-based IFFL, using Clustered Regularly Interspaced Short Palindromic Repeats (CRISPR) Cas systems for the activation arm and a viral protease for the inhibitory arm. CRISPR-Cas systems have been widely adopted due to their modularity and potential to regulate any gene. Non-mutational gene regulation systems have also been developed for targeted gene activation and knockdown, including CRISPR activation (CRISPRa) and Cas13, respectively. Lastly, these systems are relatively compact compared with many disease-relevant genes, which are simply too large to be packaged into an AAV. For example, over 300 genes that are predicted to be dosage-sensitive are longer than 4.8kb and would be difficult to deliver via AAV (**Fig 1C**).^21^ Thus, we chose to engineer CRISPR-Cas systems to act as the activation arm in our IFFL. To implement the inhibitory arm, we propose that viral proteases offer unique advantages compared to other methods. A proteolytic cleavage event does not rely on host-cell machinery, thereby improving the robustness of the system. Furthermore, this post-translational proteolytic cleavage is compatible with mRNA delivery since translation will drive both the activation and inhibition arm. Viral protease systems have also been extensively engineered for diverse controls, offering variants suitable for both *a priori* and post-delivery tuning of protease activity.^22,23^ Lastly, viral proteases have short and highly specific cleavage sites that are orthogonal to the human proteome. Thus, engineering our proteolytic IFFL simply involves introducing a proteolytic cleavage site into the CRISPR-Cas protein, thereby repressing its activity in the presence of the protease.

This proteolytic IFFL provides a novel method of modulating dosage-sensitive genes, including neurodevelopmental genes that require a narrow window of expression for healthy development. Here, we choose Retinoic Acid-Induced 1 (*RAI1*) as an example. Heterozygous loss of *RAI1* function causes Smith-Magenis syndrome (SMS), whereas its duplication causes Potocki-Lupski syndrome (PTLS).^7,9,24–26^ *RAI1* encodes a chromatin-associated transcriptional regulator that is expressed across many brain regions and in multiple cell types. ^26,27^ For example, *Rai1* in hypothalamic BDNF-producing neurons regulates mouse activity and body weight homeostasis,^28^ while complete *Rai1* loss from cortical excitatory neurons increases seizure susceptibility and dentate granule cell excitability.^29^ *RAI1* is therefore broadly acting, and correcting its dosage means restoring its expression across multiple regions and cell types.

Current curative strategies for treating SMS focus on CRISPR-mediated activation of the single remaining *Rai1* copy, as the gene is prohibitively large for AAV delivery.^30^ This preclinical CRISPRa work has shown that transcriptional upregulation of the intact allele raises *Rai1* levels and partially rescues obesity phenotypes in *Rai1*^+/-^ mice.^30^ This study intervenes in adolescence, and thus the question remains whether earlier CRISPRa intervention can rescue other phenotypes such as reduced mouse activity. In addition, they deliver CRISPRa without any dosage-control mechanism and thus inherit the delivery variability described above: *Rai1* activation in any given cell is determined by how much vector it happens to receive. High dosages risk overcorrection, pushing cells past the therapeutic range and potentially into phenotypes characteristic of the opposing disorder. Thus, a lower viral titer was used to address this concern. However, using less virus also inherently reduces the number of cells that can be treated and the potency of gene activation. Instead, using a method of controlled gene delivery would allow us to increase the bulk viral dose without the risk of overcorrection, thereby greatly improving the safety and efficacy of this strategy and making SMS an attractive application for proteolytic IFFLs.

Here, we combine experimental and computational methods to implement novel IFFLs based on proteolytic cleavage and CRISPR-mediated activation/repression. Using transient, genome-integrated, and endogenous target genes, we demonstrated the IFFLs’ feasibility, robustness, mRNA compatibility, tunability, and ability to both increase and decrease target gene expression. As a therapeutically relevant example, we applied proteolytic IFFLs to correct haploinsufficiency for SMS via CRISPRa by controlling *RAI1* activation through various delivery methods and in multiple contexts. We adapted our IFFL to target human *RAI1* and demonstrated dosage control in both HEK293 and SMS patient-derived cells. Next, we advanced the system *in vivo*, delivering the CRISPRa and proteolytic arms using a dual-AAV split-protease strategy to a phenotype-relevant brain area of wild-type (WT) mice, quantifying *Rai1* expression, and demonstrating that our IFFL produces more consistent *Rai1* activation across neurons and between animals than CRISPRa alone. Finally, we deliver the system to an SMS mouse model via neonatal intracerebroventricular (ICV) injection, intervening much earlier in disease progression than previously attempted. Alongside conventional behavioral assays, we used recently developed continuous digital home-cage monitoring system^31,32^ to track activity longitudinally, resolving phenotypes that single-session tests sample too sparsely to detect. Dosage control improved post-treatment survival and behavioral phenotype rescue in female SMS mice relative to conventional CRISPRa alone, suggesting that proteolytic IFFLs can mitigate the cost of overcorrection in a therapeutically relevant setting.

## Results

### Establishing proteolytic IFFLs for gene activation using synthetic reporters

To demonstrate the feasibility of building proteolytic IFFLs to control the dosage of CRISPRa gene activation, we first developed computational models and designed synthetic reporters to characterize circuit behavior. The activation circuit, which controls the activation of an endogenous gene of interest, consists of two components: a protease and a protease-repressible CRISPRa unit (**Fig 1D**). We reasoned that we could repress CRISPRa function by placing a protease cut site between its catalytically inactive Cas9 (dCas9) domain and the transcriptional activation domain. Thus, proteolytic cleavage would abolish the transcriptional activation ability of CRISPRa. To qualitatively explore this circuit’s behavior, we developed a computational model using Michaelis-Menten kinetics, ordinary differential equations, and parameter values in experimentally grounded regimes (**Fig 1E**). The model indeed predicted that co-delivery of CRISPRa and the protease reduces the range of target gene activation. Furthermore, increasing the protease amount relative to CRISPRa should lead to stronger dosage-control effects, thereby lowering gene expression.

Next, to experimentally validate the activation circuit, we engineered a widely used CRISPRa system, *Streptococcus pyogenes* dCas9 (SpdCas9) fused to the transcriptional activation domain VP64, to be cleavable by tobacco etch virus (TEV) protease. This was done by inserting the TEV cleavage site between the two domains. TEV protease cleaves a short, specific amino acid sequence that is orthogonal to the human proteome, avoiding off-target cleavage in the host cell. To facilitate characterization, we created a HEK293 cell line with a genomically integrated green fluorescent protein (*GFP*) reporter to act as the endogenous target gene. We then transiently transfected plasmids separately encoding all IFFL components (dCas9, TEV, and the *GFP*-targeting gRNA) and an mCherry transfection marker into the reporter cell line. Using multiple plasmids facilitates component swapping and stoichiometry titration for initial optimizations.

The first method we used to measure the IFFL’s ability to buffer against dosage variability was to transfect different amounts of IFFL plasmids and examine the average *GFP* activation, which we term “circuit titration” (**Fig 1F**). As expected, *GFP* activation in the reporter cell line was diminished upon TEV transfection compared with conditions without TEV, indicating dosage control. The second method we used to measure the IFFL’s ability to buffer against changes in dosage was to rely on the fact that plasmid uptake during transient transfection is highly correlated, meaning that mCherry fluorescence can serve as a proxy for the amount of IFFL components within a cell. This technique is widely used, and it captures a wider range of vector-level variability than circuit titration.^16,18^ Plotting the running average of GFP fluorescence against dosage (mCherry fluorescence) also confirms dosage control (**Fig 1G**). Furthermore, increasing the amount of TEV plasmid transfected decreased GFP activation, consistent with our model prediction. However, we also observed that GFP intensity decreased at high transfection levels in the presence of TEV. This trend was also observed when analyzing the running averages from the previous circuit titration experiment, suggesting that averaging across all transfected cells (**Fig S1A**) would skew the average GFP fluorescence. Thus, to extract the quantities in **Fig 1F**, we considered only cells within the monotonic regime (**Fig S1B**).

We hypothesized that this non-monotonic effect at high dosages may be due to the creation of inactive dCas9 species that bind to *GFP* and outcompete functional species. We were able to recapitulate this behavior in the simulation upon including competitive inhibition (**Fig 1H**). To further verify that this effect was indeed caused by the presence of cleaved dCas9 species, we created a “pre-cleaved” dCas9 construct that is missing the activation domain. Then we transfected both the “pre-cleaved” construct and the active CRISPRa construct into the reporter cell line. As the amount of “pre-cleaved” dCas9 increased, the average GFP expression decreased, indicating competitive binding (**Fig S1C**).

Lastly, we demonstrate the robustness of our proteolytic IFFL by comparing its performance across three different cell lines: HEK, Chinese hamster ovary (CHO), and colorectal cancer cells (COLO320DM). We genomically integrated a *GFP* reporter gene in CHO and COLO cells as we did for the HEK293 cells in Fig 1F-G. Transient transfection of the proteolytic IFFL demonstrated robust dosage control and qualitatively similar performance across all three cell lines (**Fig 1I**). This result is consistent with the fact that a protein circuit’s functionality is self-contained and principally relies on translation to function.^33^ To directly contrast it with miRNA-based IFFLs, where we expect the performance to be more variable due to dependence on the cell-specific RNAi pathways, we also tested the ability of RNAi to control the dosage of CRISPRa using a previously reported modular design^18^. As expected, the miRNA-based IFFL dosage control was more variable; furthermore, its dosage control effect was also much weaker than the proteolytic IFFL (**Fig S1D**). This may be due to the fact that a small amount of CRISPRa amplifies target gene expression, which would require substantial amounts of RNAi activity for dosage control, making the difference between these two mechanisms especially striking, whereas they might be more comparable when the target gene is directly expressed from the vector. Finally, we note that all circuits, regardless of topology, will have some cell context dependency due to the inherent problem of limited resources in mammalian cells.^19,34,35^

### Demonstrating unique features of proteolytic IFFLs

The use of proteolysis as the inhibitory arm of an IFFL enables unique features unattainable with RNA inhibition. For example, mRNA vectors have recently emerged as a safer, transient method of transgene expression. Since lipid nanoparticles carrying mRNA accumulate in the liver, potentially leading to cargo overexpression and toxicity, mRNA therapies may also benefit from dosage control offered by IFFLs.^36^ For an IFFL to function, the components constituting both arms must be constantly produced from the vector. Delivering an mRNA transcript and siRNA would alter the half-life of the mRNA rather than lead to dosage-sensitive behavior. For a proteolytic IFFL, both the CRISPR-Cas component and TEV can be encoded onto mRNAs and continuously translated by the cell. To demonstrate this, we produced *in vitro* transcribed mRNAs separately encoding the CRISPRa components, TEV, and a blue fluorescent protein (BFP) transfection marker. Since expression from mRNA is lower than DNA, we replaced the VP64 activation domain with the stronger activation domain NZF^37^ in the CRISPRa construct. Transfecting these components into the *GFP*-integrated cell line led to diminished *GFP* activation in the presence of TEV (**Fig 2A**).

**Fig 2.**
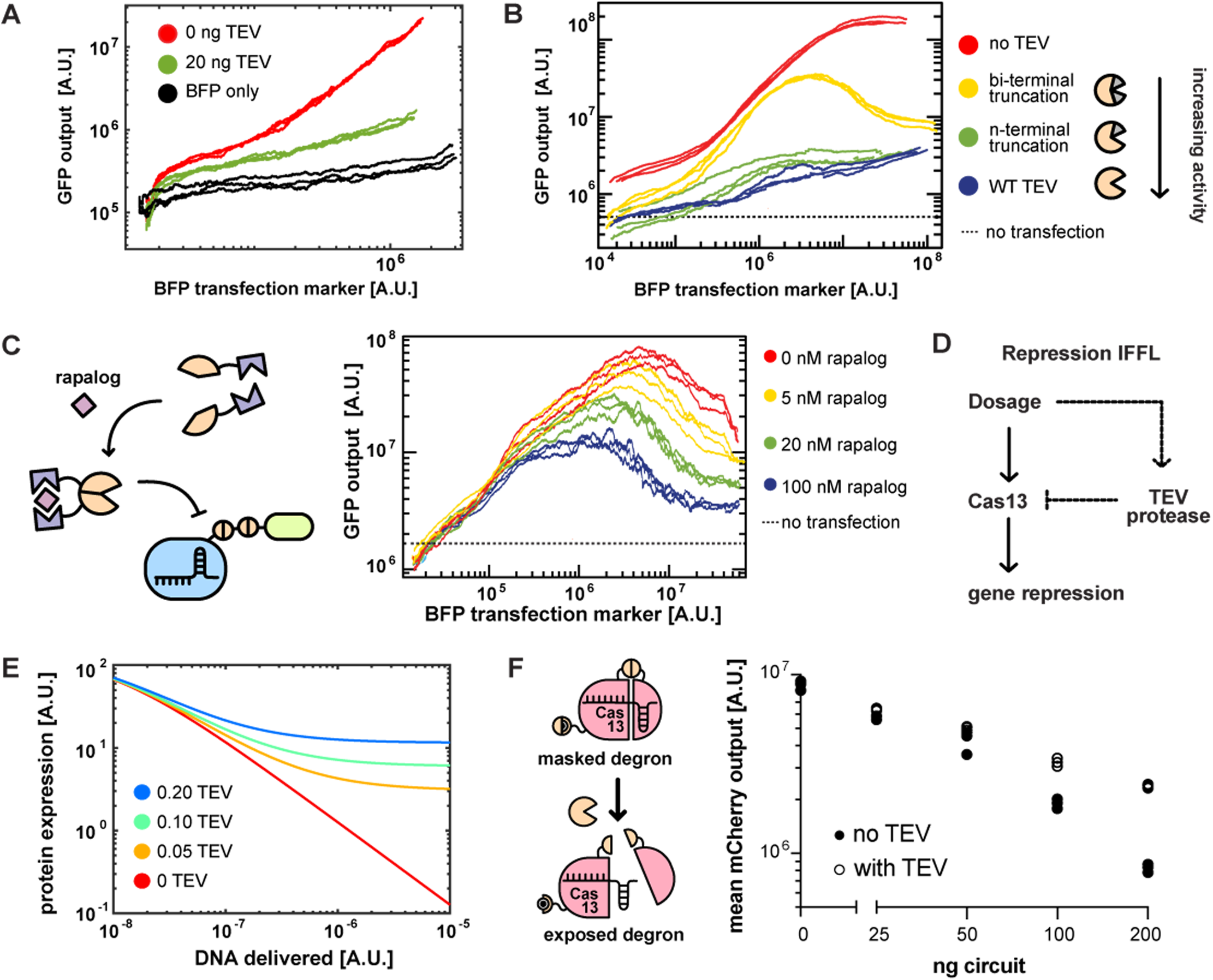
Unique features of proteolytic IFFLs. **A** Running average of GFP fluorescence (output) as a function of BFP fluorescence (transfection marker) upon transient mRNA transfection of IFFL in a GFP-integrated HEK293 cell line. **B** Running average of GFP fluorescence (output) as a function of BFP fluorescence (transfection marker) upon transient transfection of IFFL using truncated variants of TEV in a GFP-integrated HEK293 cell line. Increasing amounts of truncation reduce cleavage efficiency. **C** Schematic of the small molecule rapalog reconstituting split TEV. Running average of GFP fluorescence (output) as a function of BFP fluorescence (transfection marker) upon transient transfection of IFFL using a rapalog-induced TEV and a double TEV cut site in a GFP-integrated HEK293 cell line. **D** Schematic of engineered repression circuit. TEV protease inhibits Cas13 through proteolysis, creating an IFFL with dosage as the input and Cas13 activity as the output. **E** Simulation of the repression IFFL using ordinary differential equations. TEV fractions represent relative expression. **F** Schematic of optimized TEV-repressible Cas13 design (left). TEV inhibits CasRx by cleaving the enzyme itself and exposing a masked degron, increasing the degradation rate of CasRx. Dot plot of average mCherry fluorescence (output) as a function of circuit transfected for highly transfected cells (right). See Fig S2F for more details.

Next, we demonstrate how gene expression modulation can be achieved without varying the stoichiometry of IFFL components through *a priori* and post-delivery tuning. This is useful for viral transduction, where all components are delivered on the same construct and performance variability is minimized, but altering the strength of the repression arm is nontrivial. Therefore, to achieve *a priori* tuning of gene expression levels, we instead varied TEV cleavage efficiency to adjust the strength of the IFFL repression arm. One method to adjust cleavage efficiency is to alter the cut site. We accomplished this by inserting two adjacent TEV cut sites between dCas9 and the activation domain. This construct, co-transfected with TEV, also demonstrated dosage control but required less protease to repress gene activation (**Fig S2A**). Alternatively, we can adjust the TEV protease efficiency by truncating either the N-terminus or both termini of the protease. As the protease is truncated, it becomes less efficient, and gene expression increases even when the amount of transfected protease plasmid is kept constant (**Fig 2B, S2B**). Then, we demonstrated post-delivery tuning of gene expression using two different strategies and small molecules that control protease activity, a useful feature for adjusting gene expression level over time. First, we used the small molecule rapalog to reconstitute a split TEV protease^38^ through dimerization domains fused to each half (**Fig 2C**). Since we predicted that splitting TEV would weaken the repression arm of the IFFL, we used the dual cut site design to improve TEV sensitivity. Increasing the amount of rapalog in the media before transfection reduced gene activation. The second strategy used the FDA-approved hepatitis C virus protease (HCVP) small molecule inhibitor Voxilaprevir to tune CRISPRa activity.^39^ This involved swapping the TEV cut site between dCas9 and the transcriptional activator for the HCVP cut site. After verifying that HCVP could control the dosage of the modified CRISPRa construct (**Fig S2C**), we demonstrated that increasing the concentration of Voxilaprevir in the media increased gene activation through repressing HCVP activity (**Fig S2D**).

Finally, we designed a proteolytic IFFL to control the level of gene repression using CRISPR-Cas13 with simulations and synthetic reporters. Like the activation circuit, the repression circuit consists of two components: TEV protease and a TEV-repressible CRISPR-Cas13 (**Fig 2D**). Computational simulation suggests that such an IFFL would achieve dosage control of the down regulation of a target gene (**Fig 2E**). We chose RfxCas13d (referred to as CasRx), due to its small size and interest for therapeutics use.^40^ Since CasRx is known to have strong collateral activity that only occurs when targeting abundant mRNA transcripts,^41^ we chose a regime where collateral activity is minimized (data not shown). To establish proteolytic inhibition of Cas13, we screened for cleavage sites that minimally disrupt Cas13 activity yet render the enzyme susceptible to protease cleavage. After displaying limited inhibition effect, we added a second cleavage site at its N terminus based on the “N-end rule”^42^ such that, once exposed upon cleavage, it will serve as a degradation signal and further reduce Cas13 activity (**Fig S2E**). We then performed a circuit titration with this construct and observed dosage control for repression (**Fig 2F, S2F**).

### IFFL-mediated activation of *RAI1* in a human SMS patient cell line

After using synthetic reporters to characterize and optimize the proteolytic IFFL, we then focused on applying it to a more therapeutically relevant context: controlling the activation of the dosage-sensitive gene *RAI1* in an SMS human patient-derived cell line. First, we screened for human *RAI1*-targeting guide RNAs in HEK cells and identified one that enabled CRISPR activation of endogenous *RAI1* (**Fig S3A-D**). Next, we transfected HEK293 cells with our IFFL using this gRNA and validated that TEV represses endogenous *RAI1* activation using RT-qPCR. Results from this ensemble measurement are consistent with graded repression of CRISPRa in response to increasing TEV expression (**Fig 3A**).

**Fig 3.**
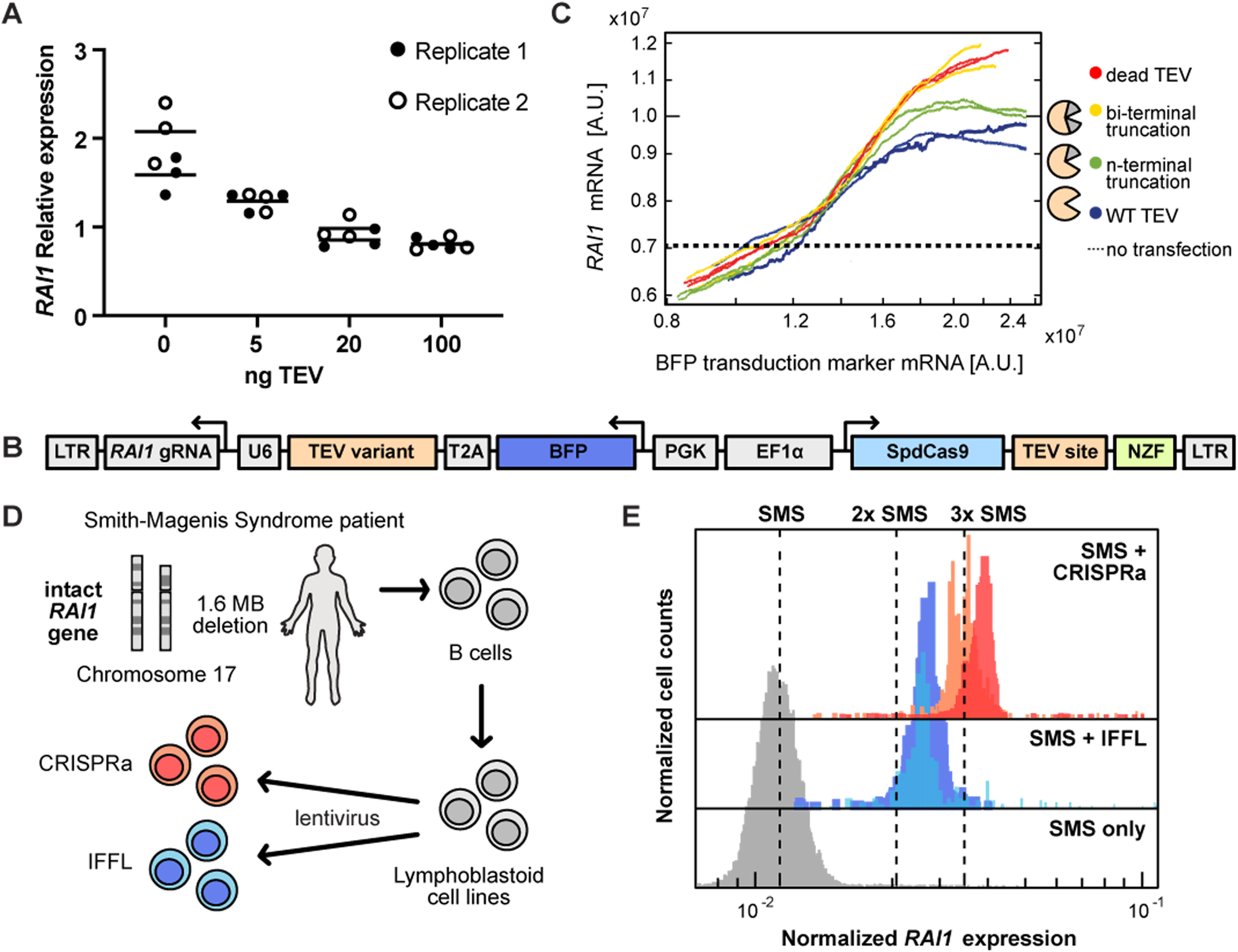
Dosage-controlled activation of *RAI1* in a Smith-Magenis patient cell line. **A** Dot plot of qPCR results using gRNA targeting *RAI1* in transiently transfected HEK293 cells. Results are based on two biological replicates (full and empty circles) and three technical replicates. **B** Schematic of IFFL lentivirus plasmid design. **C** Running medians of *RAI1* mRNA as a function of *BFP* transduction marker mRNA for lentivirus transduction of circuit in HEK293 cells. **D** Schematic of patient lymphoblastoid cell line creation and transduction. **E** Histograms of normalized *RAI1* expression for each condition of SMS LCLs transduced with CRISPRa, IFFL, or no transduction. Each color represents an independent replicate. Cells were gated on live SSC-FSC plots, high *GAPDH* expression, and positive *BFP* signal. *BFP* histograms are shown in Fig. S3E. Median levels for CRISPRa, IFFL and SMS-only histograms are 0.365, 0.276, and 0.117, respectively, averaged over the two replicates. CRISPRa and IFFL conditions have two independent replicates with a minimum of 1000 cells in each histogram. SMS-only condition has 90,000 cells. P-values were calculated based on Kruskal Wallis Test followed by post-hoc Dunn’s test. P < 1e-15 for untransduced histogram vs transduced conditions. IFFL histogram duplicates are not significantly different from each other (p = 0.60). P = 6.36e-6 for CRISPRa histogram duplicates. Finally, IFFL and CRISPRa histograms are significantly different from each other (p < 5e-04 for all comparisons).

We then proceeded to single-cell analysis. We encoded all IFFL components into a single lentiviral vector to reduce variation in circuit performance by maintaining strict stoichiometry among CRISPRa, TEV, and the gRNA (**Fig 3B**). To quantify *RAI1* expression in single cells, we adopted hybridization chain reaction (HCR) RNA-FISH to quantitatively correlate target RNA expression with fluorescence.^43^ We demonstrated that changing the efficiency of TEV encoded on the lentivirus allows us to tune *RAI1* activation in transduced HEK cells (**Fig. 3C**). Here, the condition with a catalytically dead TEV corresponds to conventional CRISPRa. We note that the activation range and tunability of *RAI1* via lentiviral transduction is much smaller than the activation of a synthetic gene during transient transfection, which is expected due to the significantly smaller variability and absolute gene copy number delivered via transduction. The relatively modest variability and nuanced dosage control here are more pertinent to the therapeutic case where there would be a critical difference between two versus three copies of a causal gene underlying a genetic disorder.

Finally, we used SMS patient-derived B-lymphoblastoid cell lines (B-LCLs), which have been used to assess the potential immune phenotype of *RAI1* haploinsufficiency, as a model system to evaluate circuit performance in a human disease-relevant SMS context (**Fig 3D**).^44^ These cells enable us to quantitatively evaluate circuit performance on a single copy of *RAI1*. Using HCR, we quantified the *RAI1* activation in the CRISPRa- and IFFL-transduced B-LCLs, compared to non-transduced LCLs. CRISPRa alone increased *RAI1* mRNA levels by almost threefold whereas the IFFL reduces activation, indicating the functionality of proteolytic dosage control (**Fig 3E, S3E**). Most notably, the CRISPRa condition exhibited a long tail of highly activated cells, whereas this tail is truncated in the IFFL conditions. This data suggest that the IFFL reduces upper-bound variability of *RAI1* activation in SMS patient-derived cell lines.

### Stereotactic AAV delivery of proteolytic IFFL to mouse brains

We then prepared to test IFFL *in vivo* to better model the physiological context that such a therapy might encounter. For a dosage-controlled *in vivo* gene therapy for SMS, which requires CNS delivery, AAVs are more suitable due to their efficiency in transducing non-dividing cells, low immunogenicity, and non-integrative gene delivery.^1,45^ To fit our IFFL onto AAVs, we designed a dual system that ensures the strength of the activation and inhibitory arms of the IFFL is coupled to the dosages of both AAVs in a cell. We replaced SpdCas9 with a shorter variant, *Staphylococcus aureus* dCas9 (SadCas9), and used the short but potent transcriptional activator NZF.^37^ The first AAV encodes SadCas9-NZF and the first half of TEV, while the second encodes a BFP transduction marker, the *Rai1* gRNA,^30^ and the second half of TEV, which is either active (IFFL) or catalytically dead (CRISPRa) (**Fig. 4A**). The split halves of TEV are fused to heterodimerization leucine zippers that reconstitute the full protease when both halves are present. Furthermore, the IFFL and CRISPRa AAVs are identical except for TEV catalytic activity (one amino acid mutation), so any differences between them are most likely due to proteolytic dosage control rather than dCas9 expression, viral load, or gRNA activity.

**Fig 4.**
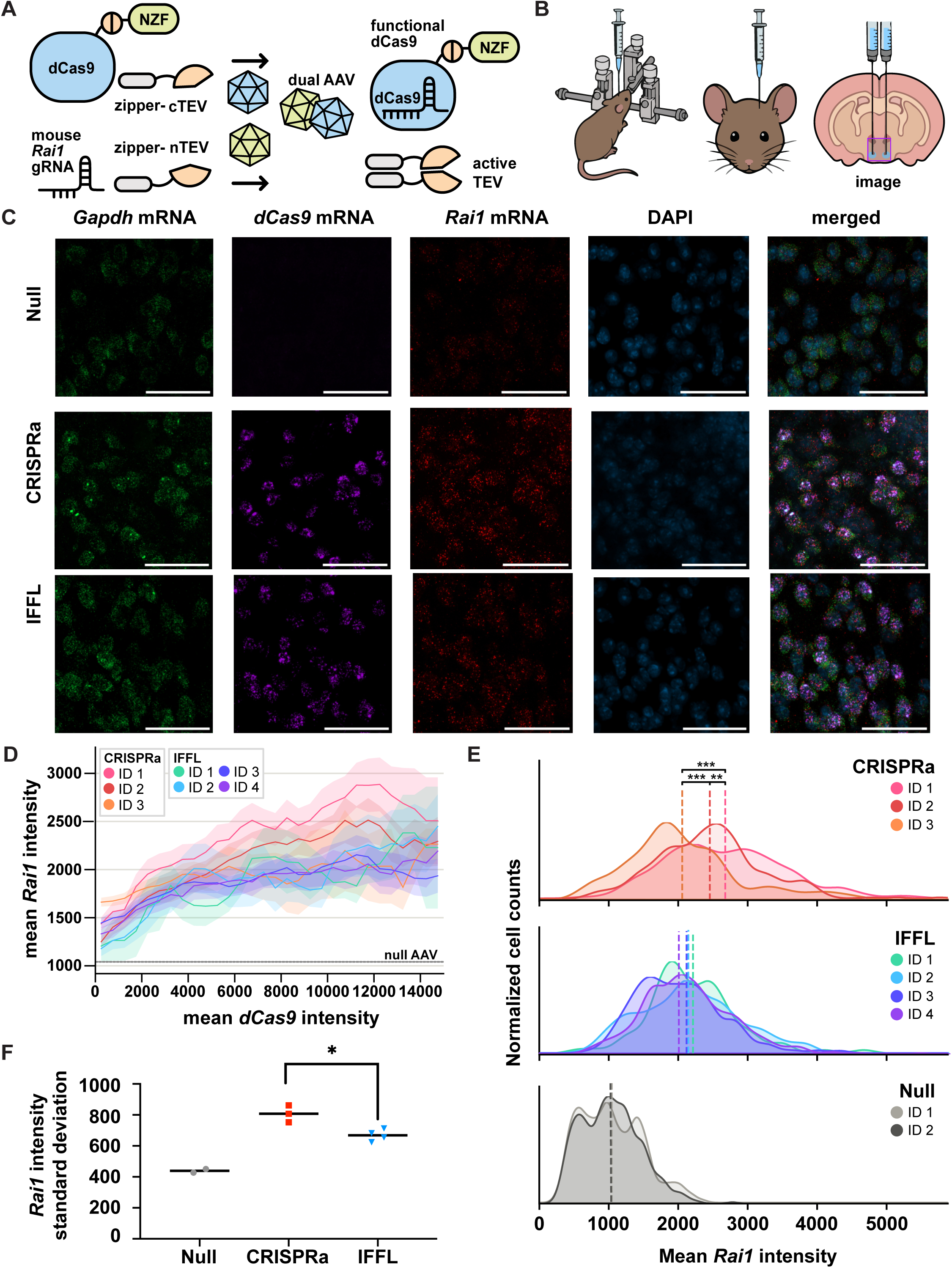
Dual AAV delivery of IFFL to adult WT mouse brains. **A** Schematic of dual AAV design. **B** Schematic of stereotactic AAV injection and location of images for Fig 4C and S4E. **C** Sample confocal microscopy images of brain slices transduced with the dual AAV system. mRNA transcripts visualized using HCR appear as bright puncta across three different channels: *Gapdh* (AlexaFluor-488), *dCas9* (AlexaFluor-546), and *Rai1* (AlexaFluor-647). Scale bar represents 50um. **D** *Rai1* mean intensity per cell and SEM as a function of binned mean *dCas9* intensity for CRISPRa (n=3) and IFFL (n=4) treated brains. Each curve consists of over 400 cells for IFFL and CRISPRa conditions. Average *Rai1* intensity for null AAV treated mice is represented by the gray line and consists of 500 cells per brain (n =2). **E** Histogram of mean *Rai1* intensity per cell gated on high *dCas9* expression (>6000) for each treated animal. Each histogram consists of 80-300 cells for IFFL and CRISPRa conditions, and over 300 cells for the null. The mean of each histogram is marked by a dashed line. P-values for difference in means calculated by the Games-Howell test for unequal variance and sample size. **F** Standard deviation of *Rai1* intensity for each histogram in Fig 4E. Welch t-test was used to calculate p-value for unequal variance.

We first validated that this dual AAV design can control *Rai1* activation *in vitro* using N2A cells (a mouse neuroblastoma cell line) and then in primary mouse cortical neurons. We used HCR and flow cytometry to quantify single-cell *Rai1* activation in transduced N2A cells, observing that the IFFL system has lower normalized *Rai1* activation than CRISPRa alone and a lack of highly activated cells (**Fig S4A-B**), consistent with our observations in LCLs. Then, we used HCR and confocal microscopy to visualize mRNA transcripts in transduced primary mouse cortical neurons and quantified the *Rai1* expression by calculating the percent area per cell that is positive for *Rai1* (**Fig S4C-D**). The *Rai1* histograms for the CRISPRa and IFFL conditions were qualitatively consistent with our other *in vitro* experiments, indicating that the dual AAV IFFL system reduces upper-bound variability of *Rai1* activation *in vitro*.

Next, we administered the dual AAV system to 8-week-old WT mice via stereotactic injection to the paraventricular nucleus of the hypothalamus (PVH) (**Fig 4B**). This region is of therapeutic relevance to SMS: a previous study demonstrated that activation of *Rai1* via CRISPRa in the PVH of 4-week-old *Rai1*^+/-^ mice partially rescued obesity phenotypes.^30^ Thus, we chose to use this region for quantitative in vivo characterization. After four weeks, the brains were harvested, and HCR was used to probe for *Gapdh* (housekeeping gene), *dCas9*, and *Rai1* mRNA transcripts, which were then captured using confocal microscopy (**Fig. 4C**). As with the *in vitro* data, there are visually more *Rai1* transcripts in the CRISPRa and IFFL conditions in *dCas9*-positive cells compared to a null AAV. A tiled image of the *dCas9* signal captures the dosage gradient, with the brightest signal aligning with the injection site and declining as one moves farther away (**Fig S4E**). To quantify the correlation between *Rai1* and *dCas9* signals, we used an AI-based pipeline to calculate the mean intensity per cell for over 4000 cells across multiple brain slices and animals (**Fig 4D**). As *dCas9* signal increases, *Rai1* intensity appears to increase more for the CRISPRa conditions compared to IFFL, which slightly plateaus. Furthermore, the spread of the *Rai1* signal at any dCas9 intensity appears larger with CRISPRa than with IFFL. To quantify the variability of *Rai1* activation, we plotted histograms of mean *Rai1* intensity for high expressing *dCas9* cells for each treated animal (**Fig 4E**). The CRISPRa-treated brains show significantly different histograms of *Rai1* intensities, indicating three distinct underlying distributions, whereas the difference for IFFL is not significant, making them statistically identical. The standard deviation of the *Rai1* intensity histograms is also significantly higher for CRISPRa than IFFL (**Fig 4F**). Taken together, this indicates that compared to CRISPRa alone, our IFFL circuit leads to more consistent gene activation across cells and across animals.

### Dosage-controlled activation of *Rai1* rescues multiple SMS phenotypes in mice

Finally, we turn to *Rai1* heterozygous mice to evaluate whether dosage control can rescue disease phenotypes and improve outcomes compared to CRISPRa alone in a preclinical model of SMS. *Rai1*^+/-^ mice recapitulate selective core features of human SMS, including obesity and behavioral and activity deficits, making it a tractable system for testing dosage-correcting therapies before clinical translation.^26,46,47^ Phenotype severity is sex-dependent, with female *Rai1*^+/−^ mice showing a more penetrant obesity phenotype than males.^47^ Unless otherwise noted, we focus on female cohorts throughout this section. Although most male results (Fig. S5-S6) don’t contradict the trends we observed in females, there is often no significant difference between SMS-IFFL and SMS-CRISPRa males, suggesting that the male experiments might be underpowered or the dosage effects might be sex-specific.

*Rai1* is broadly expressed and developmentally timed, rising through late embryogenesis, peaking approximately one week after birth, and remaining high throughout postnatal life.^26^ Temporal dissection distinguishes the consequences of *Rai1* deficit at different developmental stages,^26,48^ indicating that the neurobehavioral phenotype depends on *Rai1* function during a developmental window that has closed by three weeks of age, whereas the metabolic phenotype reflects a requirement that persists into adulthood. Effective intervention must therefore be early, before the neurobehavioral window closes, and broad, since no single brain region accounts for the full phenotype. Neonatal intracerebroventricular (ICV) injection satisfies both requirements, being administered soon after birth and distributed throughout the ventricular system and adjacent parenchyma without requiring region-specific targeting.^49^ Thus, bilateral injections were performed into the lateral ventricles of P0-P2 pups. As controls, both WT and SMS pups received a GFP-expressing null AAV (WT-null, SMS-null) while the CRISPRa and IFFL dual AAV systems were administered to SMS pups only (SMS-CRISPRa, SMS-IFFL). Mice were monitored longitudinally for survival, body weight (4 weeks onwards), conventional behavioral tests (6 weeks onwards), and home-cage activity by Digital Ventilated Cage (DVC) analytics (20 weeks onwards) (**Fig 5A**).

**Fig 5.**
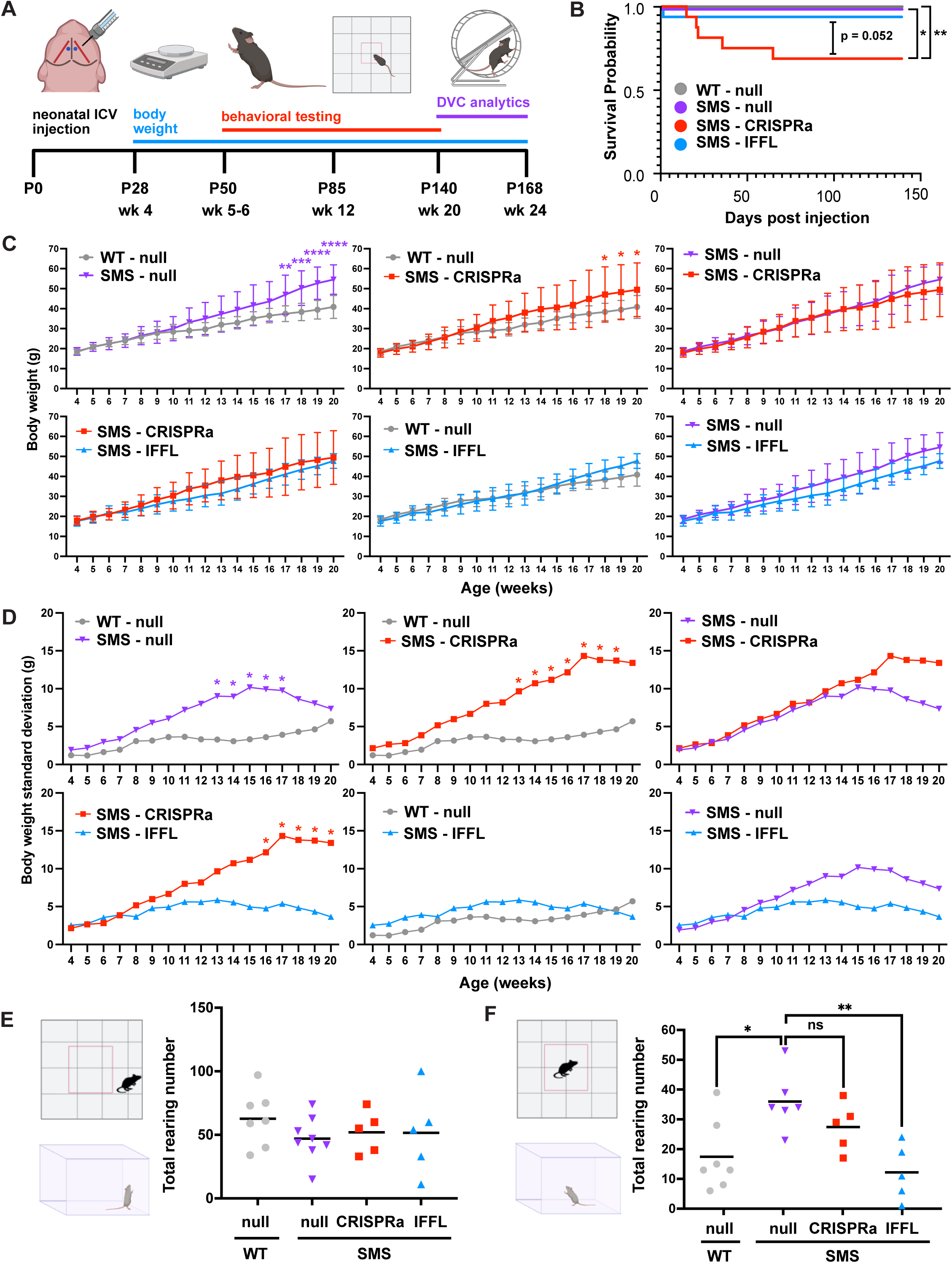
Phenotypic assessment of IFFL/CRISPRa-treated SMS mice. **A** Schematic of AAV treatment and phenotype tests. **B** Survival curves for each group (male and female combined). Number of mice is as follows: WT-null= 24, SMS-null = 16, SMS-CRISPRa = 16, SMS-IFFL = 19. Log-rank test for Kaplan-Meier curves was used to calculate p-value (uncorrected for multiple comparisons). **C** Body weight comparisons for female mice in each group ages 4-20 weeks. Two-way ANOVA followed by a Tukey’s multiple comparisons test was used to calculate p-value. **D** Standard deviation of body weight for female mice in each group ages 4-20 weeks. Levene’s test for equality of variances was used to calculate p-value. **E** Total rearing number in the periphery of the open field for female mice in each group. Ordinary one-way ANOVA followed by Tukey’s multiple comparisons test was used to calculate p-value (all comparisons found to be not significant). **F** Total rearing number in the center of the open field for female mice in each group. Ordinary one-way ANOVA followed by Tukey’s multiple comparisons test was used to calculate p-value.

Survival after neonatal ICV injection is generally not affected by the injection itself and instead largely depends on minimal disruption to maternal care to avoid rejection in the days after surgery.^49^ Accordingly, WT-null and SMS-null control mice achieved 100% survival throughout the study. SMS-IFFL mice survived at rates comparable to WT-null and SMS-null controls (94%), with a single death occurring at day three, within the sensitive period for maternal rejection post-surgery. In contrast, SMS-CRISPRa mice showed significantly reduced survival (69%) compared to null AAV groups, with deaths occurring throughout the monitoring period (**Fig 5B**). This suggests that uncontrolled and developmentally early *Rai1* activation may be toxic *in vivo* and that proteolytic dosage control may mitigate this toxicity. Toxicity from overexpression of a dosage-sensitive gene has been reported before in studies treating a mouse model of Rett syndrome (*Mecp2* haploinsufficiency) using neonatal ICV injection of an AAV expressing *Mecp2* without dosage control, lending support to this possibility.^20,50^

Body weight trajectories of female WT-null and SMS-null mice show a diverging trend with SMS mice becoming heavier after 10 weeks of age, consistent with the overweight phenotype associated with SMS (**Fig 5C**).^26,28,30^ We also monitored SMS-CRISPRa and SMS-IFFL mice for weight gain over the course of the experiment, and calculated animal-to-animal variability within each condition (**Fig 5D**). SMS-CRISPRa mice show significantly higher variability than SMS-IFFL and WT-null groups after week 16 and week 13, respectively. On the other hand, SMS-IFFL and WT-null weight variability remained comparable throughout. SMS-null weight variability was higher than WT-null and SMS-IFFL, but less than SMS-CRISPRa. This suggests that proteolytic dosage control constrains animal-to-animal variability in body weight, while CRISPRa produces increasingly heterogeneous, less predictable weight outcomes. This result is consistent with our quantitative understanding of how these two circuits behave (**Fig 4**): CRISPRa leads to more variable *Rai1* activation and possibly more variable phenotype than IFFL. Perhaps due to such variability, it could not be concluded whether the mean body weights differ between the SMS-IFFL and SMS-CRISPRa groups with statistical significance.

We also performed a suite of standard behavioral analysis in an open-field test in a manual, blinded fashion: total distance traveled (no significant difference, not shown), time spent in different regions of an arena (no significant difference, not shown), and repetitive rearing, when the mouse stands upright on its hind legs.^30^ We found that rearing in the periphery did not differ across groups for female mice (**Fig 5E**), whereas in the center of the field, female SMS-null mice showed significantly more repetitive rearing than WT-null controls. IFFL treatment reduced center rearing to control levels, whereas SMS-CRISPRa mouse rearing was not significantly different from SMS-null (**Fig 5F**). While statistical ambiguity remains, this provides initial evidence that there may be a behavioral difference between the SMS-IFFL and SMS-CRISPRa groups.

While the open-field environment captures locomotor behavior for a short duration in an environment novel to the mice, DVC analytics offer a method to automatically monitor activity within the home cage over long time scales with minimal human intervention.^31,32^ It has been used to characterize circadian phenotypes, such as rest disturbances in a mouse model of ALS.^51^ SMS mice are known to have greater light sensitivity than WT mice and suppressed locomotor activity during the night.^52,53^ Therefore, we decided to further analyze our cohorts using DVC analytics. The mice were housed in DVC cages for 4 weeks starting at week 20. These cages consist of a floor with movement sensors and a running wheel for continuous electrode-based activity recordings (**Fig 6A**). Individual daily floor activities were resolved and averaged across all individuals within each group (**Fig 6B-C**). Female WT-null and SMS-null mice show stark activity differences during the dark phase, with SMS-mice clearly showing lower relative activity. Notably, the SMS-IFFL but not SMS-CRISPRa group shows more activity during the night than SMS-null.

**Fig 6.**
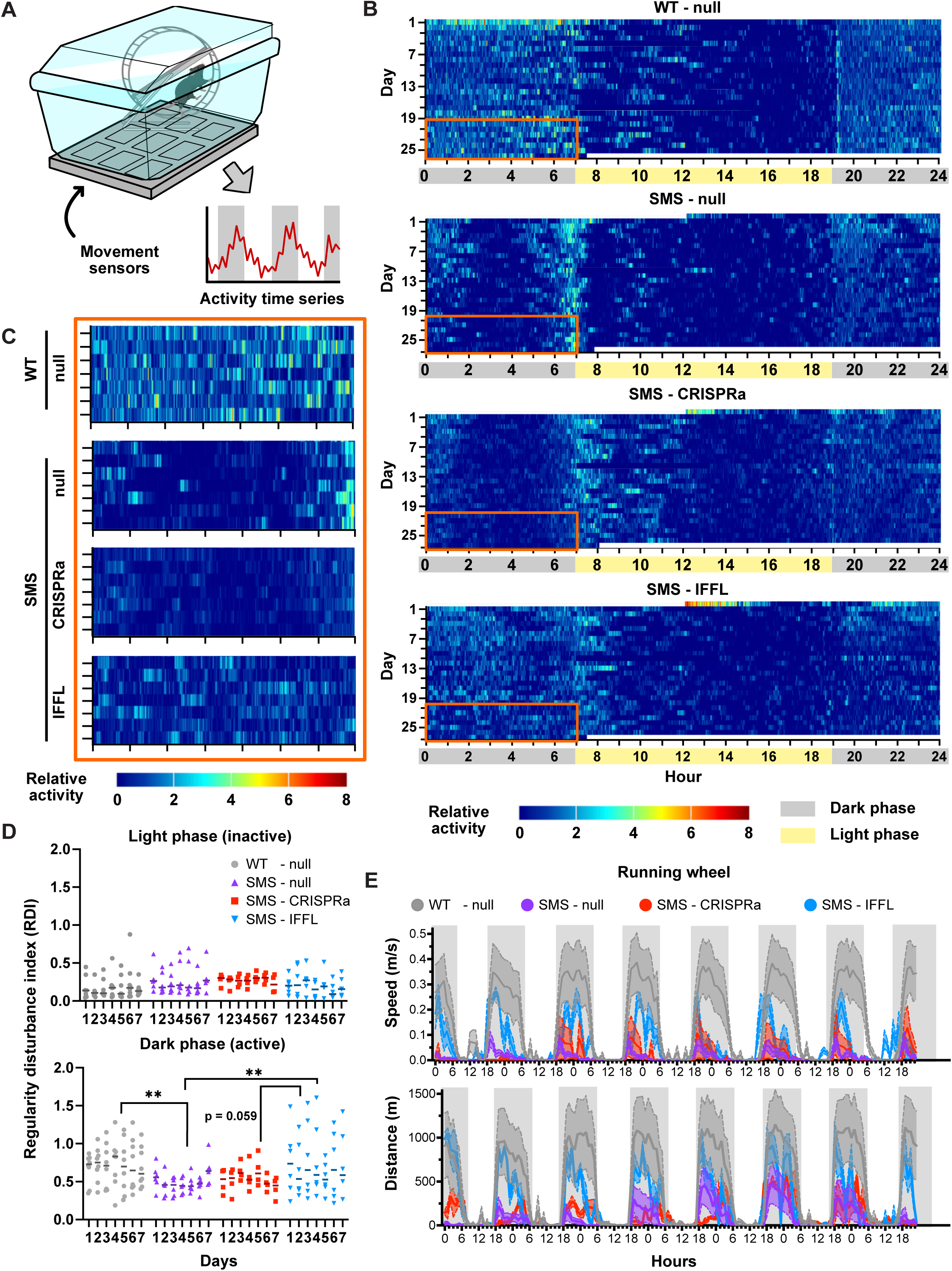
DVC and activity analytics of IFFL/CRISPRa-treated SMS mice. **A** Schematic of DVC cage with movement sensors and running wheel. **B** Heat maps of average relative activity over four weeks for female mice in each group (WT-null, SMS-null, SMS-CRISPRa, SMS-IFFL). Representative dark phase activity during the last week when the mice are most acclimated is boxed in orange (see Fig6C). **C** Heat maps of representative dark phase activity of female mice in each group during the last week of DVC data collection (boxed area in Fig 6B). **D** Dot plot of light phase and dark phase regularity disturbance index of female mice in each group. Two-way ANOVA followed by Tukey’s multiple comparisons test was used to calculate p-value. **E** Mean and SEM of running wheel speed and distance of female mice in each group.

Using this activity data, we can also calculate a regularity disturbance index (RDI), which has been previously used as a digital biomarker for quantifying progression of ALS in mice.^51^ RDI is a measure of day-to-day variability/unpredictability in activity patterns. We likewise calculated RDI for all groups and both phases. In the light phase, when animals are normally at rest, RDI did not differ substantially across groups for both female and male mice, consistent with low activity levels leaving little room for group differences to emerge. However, in the dark phase, SMS-null mice showed significantly lower RDI than WT-null mice for the female cohort (**Fig. 6D**). This reflects more predictable hypoactivity, consistent with the increased repetitive behavior associated with SMS. For female mice, IFFL treatment significantly increased dark-phase RDI relative to SMS-null, shifting activity variability toward the control pattern, whereas SMS-CRISPRa mice showed RDI levels that remained low and did not differ significantly from SMS-null mice.

As an independent measure of active-phase activity, running-wheel speed and distance were also recorded in the home cage over successive light-dark cycles (**Fig 6E**). Female SMS-null mice are distinguished from WT-null by their low running wheel speed and shorter distance covered, consistent with the literature.^53^ The speed and distance of female SMS-IFFL mice fell between the two groups, indicating partial restoration of active-phase activity, whereas female SMS-CRISPRa mice remained closer to SMS-null, indicating limited rescue.

Together, these results demonstrate the viability of using neonatal ICV injection to intervene much earlier in the development of SMS to rescue several phenotypes. Especially in females, the IFFL reduced weight variability across animals and rescued SMS phenotypes across multiple metrics–post-treatment survival, repetitive rearing, and active-phase activity–more effectively than CRISPRa alone. Thus, these results indicate that proteolytic dosage control confers an organism-level therapeutic advantage over conventional CRISPRa.

## Discussion

Dosage control via IFFLs has long been recognized as important for improving gene-manipulation techniques in basic research and for biomedical applications sensitive to gene dosage. Most implementations of an IFFL rely on delivering an exogenous gene inhibited by a self-targeting miRNA.^12,16,18,20^ Here, we introduce a new modular implementation of the IFFL that relies on proteolytic cleavage to deactivate an engineered CRISPR-Cas system. We first characterized this design and established its feasibility using simulations for guidance. Using a synthetic reporter, we observed dosage control of CRISPRa spanning over two orders of magnitude with relatively low amounts of TEV. Our simulation was particularly insightful for understanding the role of competitive inhibition, which can explain the diminishing activation towards very high transfection amounts. We then demonstrated how our protease-based IFFL tackles a few fundamental limitations of previous IFFL systems. We established the robustness of circuit performance across diverse cell types, compatibility with mRNA delivery, and the ability to tune gene expression pre- and post-delivery. Using CRISPR-Cas13, we achieved dosage-controlled gene repression, which has not been demonstrated previously.

We then applied the circuit to control the activation of the dosage-sensitive gene *RAI1* as a therapeutically relevant example of our proteolytic IFFL’s capabilities. *RAI1* haploinsufficiency and triplosensitivity lead to reciprocal neurodevelopmental disorders, SMS and PTLS, respectively.^7,9,24–26^ Thus, targeting *RAI1* via CRISPRa in an SMS patient carries the risk of overcorrection. Furthermore, patient-to-patient variability leads to unpredictable differences in gene therapy outcomes. We demonstrate how an IFFL can mitigate both these concerns using patient-derived cells and a mouse SMS model. First, we demonstrated how transduction of our circuit into a human SMS patient cell line can both more moderately activate *RAI1* than CRISPRa alone and reduce the number of highly expressing cells. Next, we designed a dual AAV system that couples the strength of the activation and inhibitory arms of the IFFL to the dosage of both AAVs in a cell for *in vivo* delivery. Neither dCas9 nor the reconstituted protease is functional unless a cell receives both vectors, so no subpopulation carries unbuffered CRISPRa. We demonstrate that stereotactic AAV delivery of the IFFL to the PVH region of the brain of adult WT mice produces more moderate and less variable gene activation across transduced cells as well as more uniform activation across animals.

Lastly, we demonstrated that our IFFL improves survival and rescues disease phenotypes more effectively than CRISPRa when delivered via neonatal ICV injection in SMS mouse models. The death of CRISPRa-treated mice within weeks following injection was particularly notable, especially since the only difference between the payload of the CRISPRa and IFFL dual AAV systems was a single amino acid mutation in the TEV protease. This leads us to speculate that the CRISPRa deaths were due to overactivation of *Rai1* rather than any potential toxicity from the AAV itself. Similar toxicity findings have been reported when overexpressing the dosage-sensitive gene *Mecp2* in a mouse model of Rett syndrome (*Mecp2* haploinsufficiency), specifically from high levels of the protein.^20,50^ Mean body weight trajectories were unchanged by either treatment relative to SMS-null, but animal-to-animal variability was several-fold lower under the IFFL and comparable to wild-type, mirroring the narrower activation distribution measured after stereotactic delivery. Rescue of disease phenotypes such as repetitive rearing, reduced activity, and reduced RDI were at least partial in the IFFL groups. Intriguingly, the CRISPRa group often had no significant difference with the SMS-null control group or worse rescue than the IFFL, especially in the female cohort. Given that we expect *Rai1* expression to be higher in the CRISPRa group than IFFL, we would predict either fuller correction than the IFFL or overshoot toward PTLS-like phenotypes, and neither was observed. One potential explanation for this incongruency may be that our mice with the highest *Rai1* activation were the ones that died. Thus, our CRISPRa cohort that reached the behavioral study stage was biased towards mice on the lower end of *Rai1* activation. Alternatively, it is also possible that high Rai1 overexpression in a heterogeneous subset of neurons leads to different phenotypes than global triplosensitivity in PTLS. Nonetheless, we were able to validate the feasibility of early intervention for SMS via ICV, introduced the use of DVC cages as a tool and RDI as a metric for characterizing SMS behavior, and demonstrated that proteolytic IFFLs confers a therapeutic advantage over unbuffered CRISPRa, principally by widening the margin between an effective and a harmful dose.

## Limitations of the study

Though we introduced a new implementation of the IFFL that relies on proteolysis and CRISPR-Cas systems and demonstrated its ability to improve gene therapy outcomes for one dosage-sensitive disease, SMS, there are still a few caveats to consider and opportunities for future work. First, at high transfection levels, we observed a non-monotonic effect in gene activation, which is not characteristic of an IFFL, which we attribute to competitive inhibition at high levels of the IFFL components achieved with transient transfection (**Fig. 1H, Fig. S1C**). Notably, this behavior is not observed for more therapeutically relevant viral delivery methods where delivered doses are lower. Secondly, further work could be done to advance the repression circuit. For example, high-fidelity versions of CasRx can be adapted for an IFFL to reduce the amount of off-target mRNA cleavage.^54^ Characterization of the repression IFFL would also benefit from targeting an endogenous transcript, such as *RAI1*. Another caveat to consider is that the human B LCLs are not the most relevant human *in vitro* model of SMS, especially since SMS is a neurodevelopmental disorder that requires correction of *RAI1* expression in the brain. Furthermore, measuring *RAI1* expression using HCR on B LCLs from SMS, PTLS, and healthy patients shows significant overlapping *RAI1* expression, making it less clear what the boundary is between SMS, healthy, and PTLS phenotypes (**Fig S3F**). Future work could implement a proteolytic IFFL in SMS patient iPSC-derived cortical neuron models or brain organoids. Lastly, our proteolytic IFFL relies on bacterial and viral proteins which may pose an immunogenic risk. It is to our advantage to target the brain, as it is an immune-privileged organ, thereby mitigating risks. This is further substantiated by the low mortality in the IFFL-treated group, indicating that toxic *Rai1* activation, rather than the AAVs or protein components themselves, caused the CRISPRa-related deaths. Nonetheless, further work would be required to determine the cause of death in these animals. Finally, there are multiple studies by us and others that explore engineering human proteases or protein deimmunization as less immunogenic alternatives.^55,56^

---

**Fig S1.**
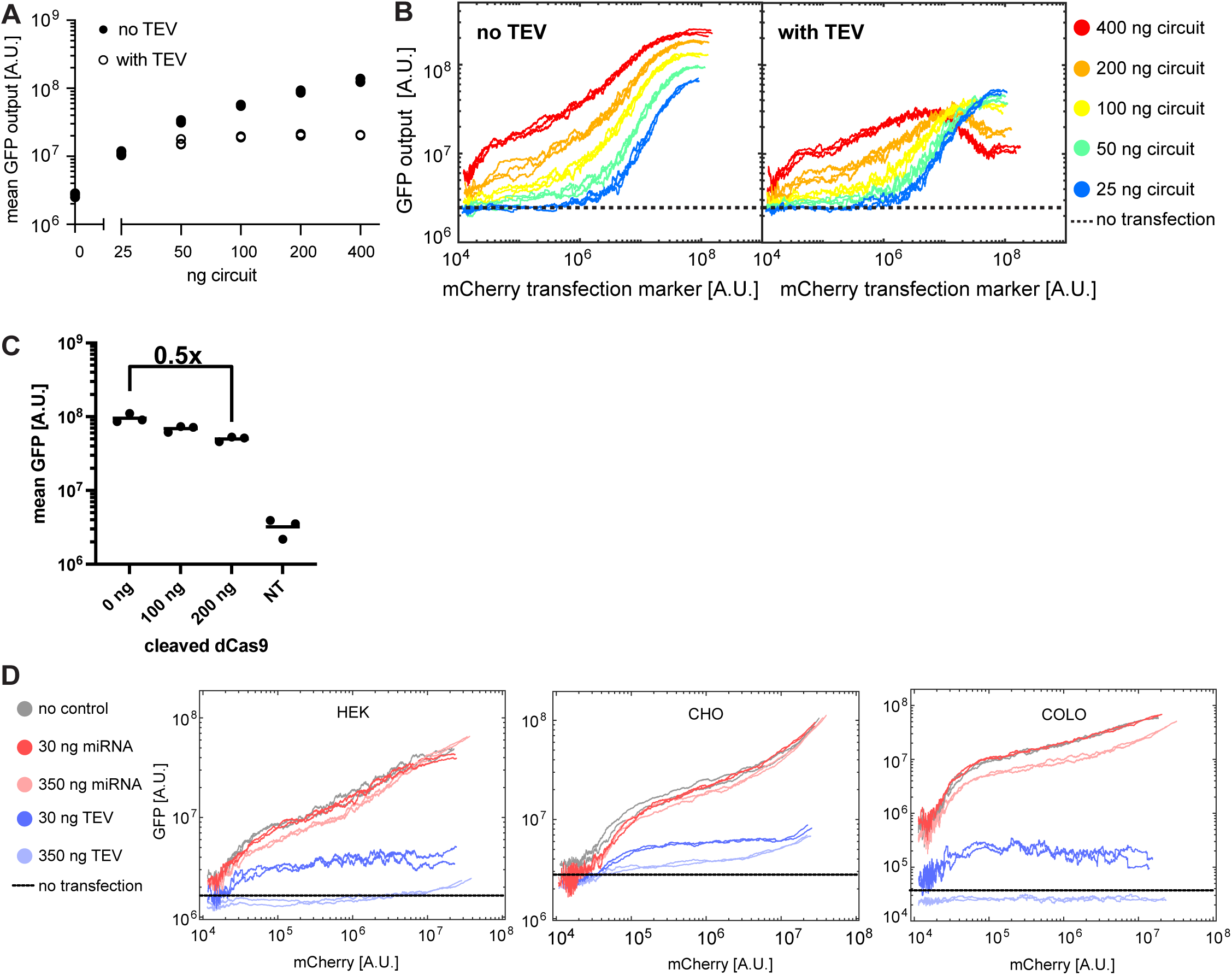
**A** Dot plot of mean GFP (output) of all transfected cells as a function of circuit transfected in GFP-integrated HEK293 cells. **B** Running means of GFP (output) as a function of mCherry transfection marker for activation circuit titration. In highly transfected cells, there is a non-monotonic effect where GFP decreases. **C** Dot plot of mean GFP expression as more “pre-cleaved” dCas9 species are transfected along with a constant amount of active dCas9 species. NT = no transfection (untransfected GFP fluorescence). **D** Running average of GFP fluorescence (output) as a function of mCherry fluorescence (transfection marker) upon transient transfection of proteolytic activation IFFL and miRNA IFFL across three GFP-integrated cell lines.

**Fig S2.**
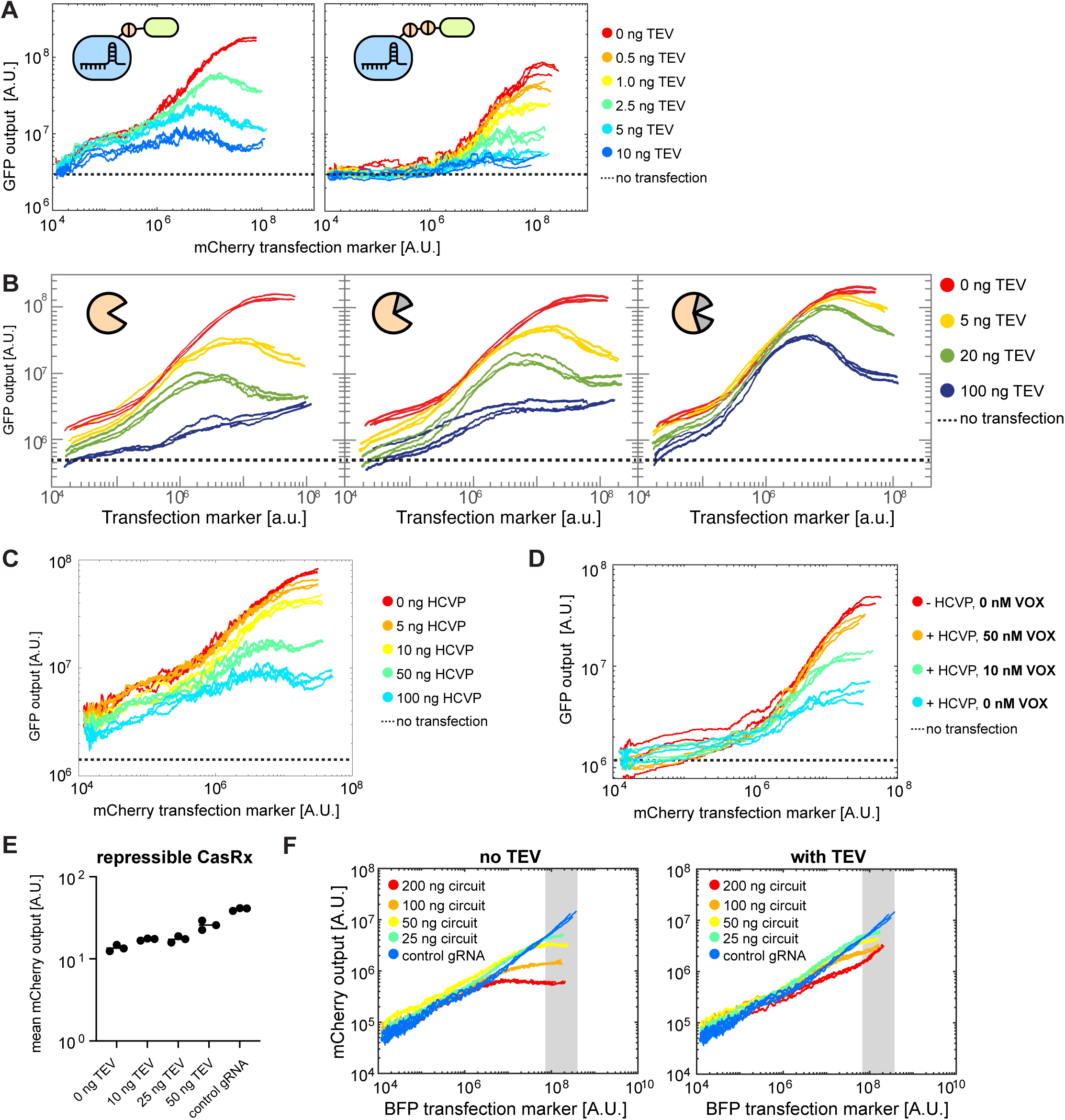
**A** Running average of GFP fluorescence (output) as a function of mCherry fluorescence (transfection marker) upon transient transfection of IFFL with a single (left) or double (right) TEV cut site in a GFP-integrated HEK293 cell line. **B** Running average of GFP fluorescence (output) as a function of BFP fluorescence (transfection marker) upon transient transfection of IFFL different amounts of TEV variants in a GFP-integrated HEK293 cell line. **C** Running average of GFP fluorescence (output) as a function of mCherry fluorescence (transfection marker) upon transient transfection of IFFL with HCVP and HCVP cut site in a GFP-integrated HEK293 cell line. **D** Running average of GFP fluorescence (output) as a function of mCherry fluorescence (transfection marker) upon transient transfection of IFFL using HCVP and Voxilaprevir (HCVP inhibitor) to tune GFP activation in a GFP-integrated HEK293 cell line. **E** Dot plot of optimized cleavable CasRx ability to repress expression of a co-transfected mCherry target plasmid as the amount of co-transfected TEV plasmid increases. **F** Running averages of mCherry fluorescence (output) as a function of BFP transfection marker for whole circuit titration experiment (Fig 2F). Gray area is the gate for highly transfected cells used to create Fig 2F.

**Fig S3.**
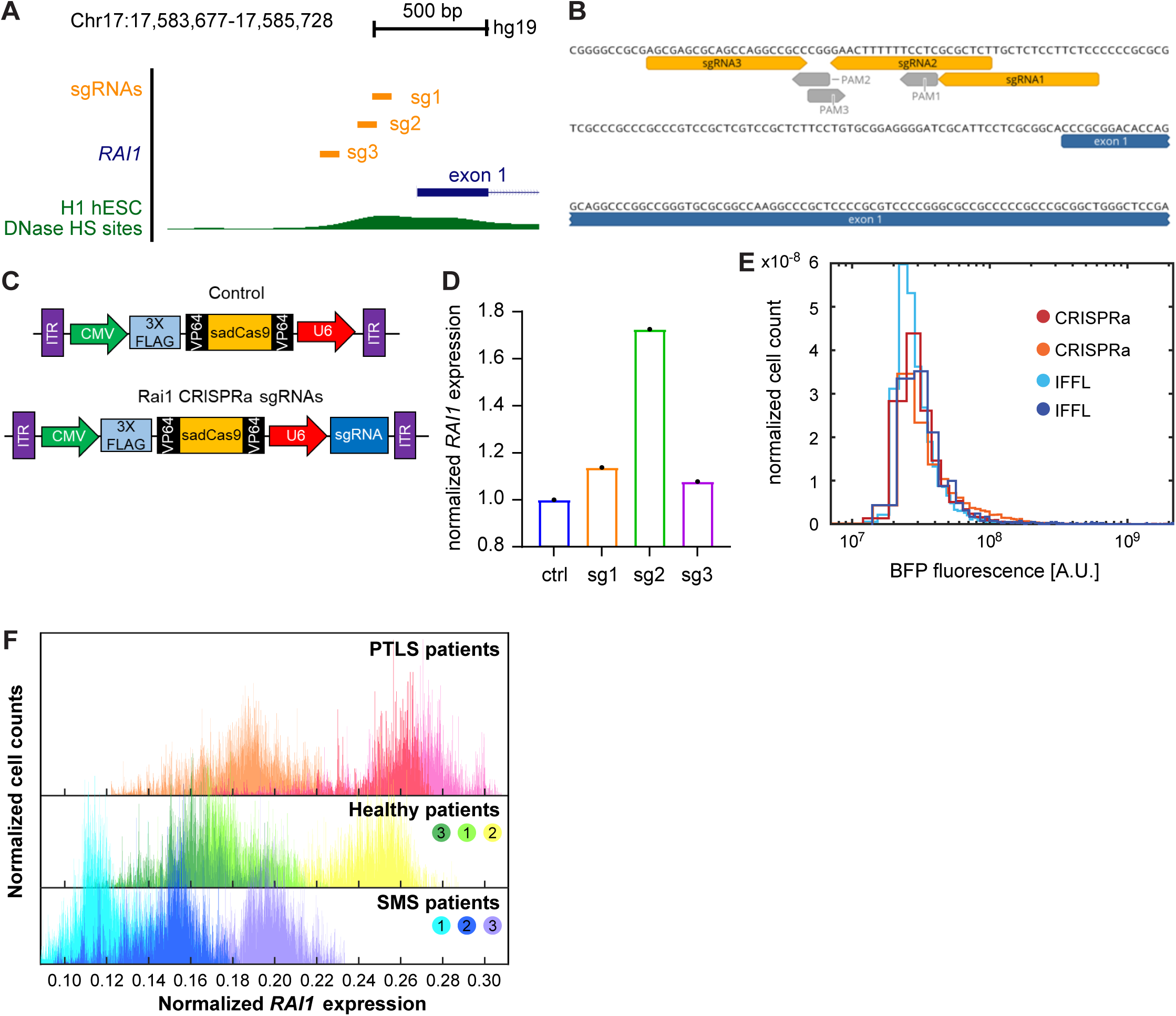
**A I**dentification of a human *RAI1* sgRNA that upregulates *RAI1* expression in 293A cell line. Schematic representation of the positions of sgRNAs (orange) in the human *RAI1* promoter. DNase hypersensitive sites (HS) in H1 human embryonic stem cells with open chromatin configurations are shown in green. **B** Positions and sequences of three candidate sgRNAs (orange) followed by Sa protospacer adjacent motif (PAM) sequences (gray) within human *RAI1*’s proximal promoter. **C** Schematic representation of the sadCas9-2 × VP64 constructs used in gRNA screen. **D** Bar plot of qRT-PCR results after sadCas9-2 × VP64-sg2 vector transfection in HEK293 cells. **E** Histograms of *BFP* fluorescence for SMS patient B-cells transduced with either CRISPRa or the feedforward circuit. **F** Histograms of normalized *RAI1* expression (*RAI1*/*GAPDH* fluorescence) for different SMS, healthy, and PTLS patient B LCL cell lines (n=3 for each group). Healthy patients are the parents of the SMS patients. Related patients are marked with the same number (1-3).

**Fig S4.**
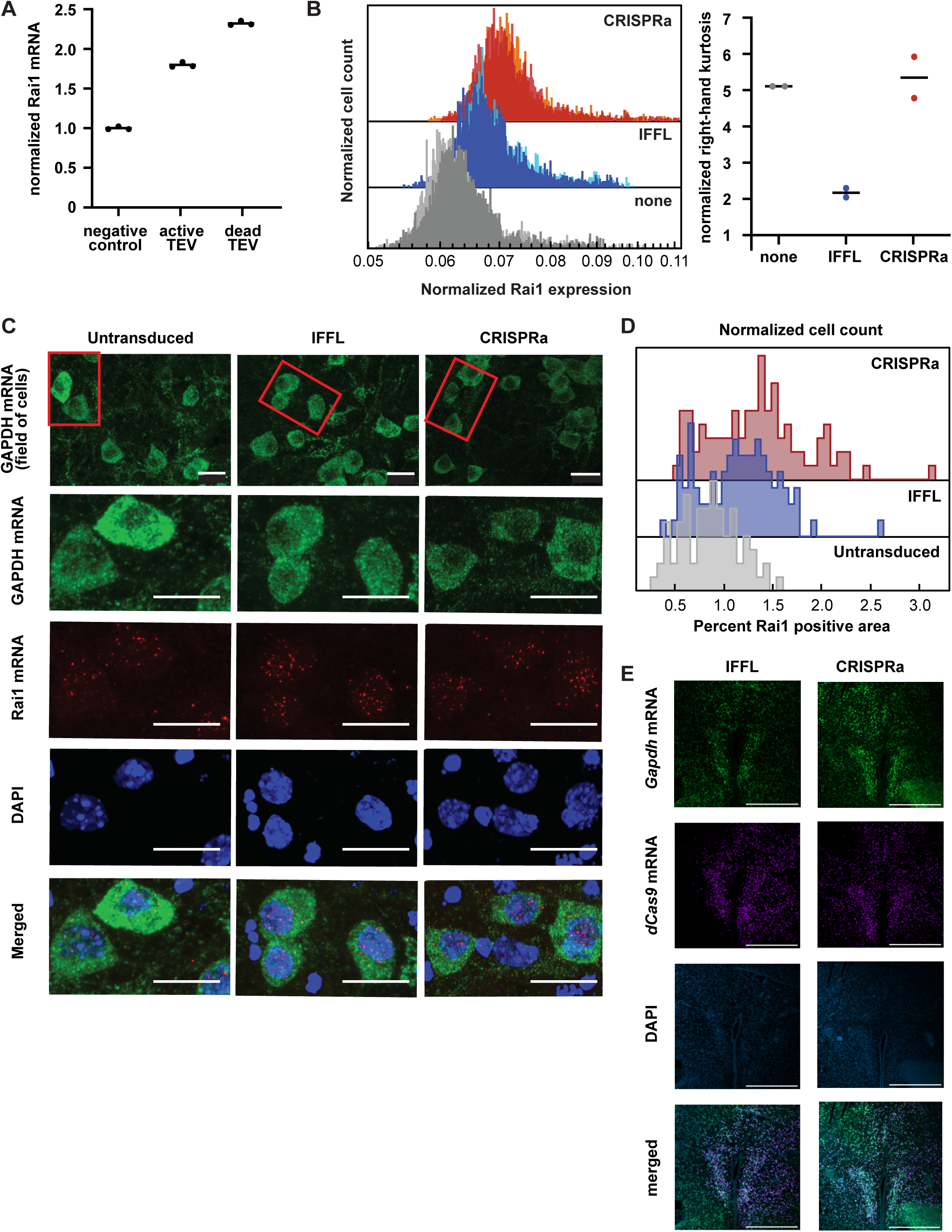
**A** Dot plot of qPCR results of N2A cells transiently transfected with two plasmids encoding the dual AAV system. **B** Histograms and corresponding right hand kurtosis of running medians for normalized Rai1 expression in N2A cells transduced with the dual AAV system containing either CRISPRa or IFFL conditions. Cells were gated on positive-stained cells, FSC-SSC live cells, with high GAPDH expression, and positive BFP signal. All conditions had two technical replicates with at least 25,000 cells in each histogram. **C** Confocal microscopy images of primary mouse cortical neurons transduced with the dual AAV system. *Gapdh* (AlexaFluor-488) and *Rai1* (AlexaFluor-647) transcripts were measured using HCR with DAPI stain in the mounting reagent. Red boxes indicate the field of view of zoomed-in images. The range for all red channel images was uniformly adjusted to better display *Rai1* mRNA puncta. Scale bars represent 20um. **D** Histograms depicting the percentage of *Rai1* positive pixels out of total soma region of individual neurons. Data for all three experiments was combined from two independently transduced wells. The untransduced histogram contains ∼80 cells, and both CRISPRa and IFFL ∼100 cells. Median percentages were increased by 35% (IFFL) and 58% (CRISPRa) relative to untransduced. P-values were calculated using Kruskal Wallis Test followed by post-hoc Dunn’s test: 5.64e-6 for untransduced vs IFFL, 5.44e-12 for untransduced vs CRISPRa, and 1.68e-2 for IFFL vs CRISPRa. **E** Representative and tiled confocal microscopy images of the injection site for IFFL and CRISPRa treated brain slices. *Gapdh* transcripts (AlexaFluor-488) and *dCas9* transcripts (AlexaFluor-546) were visualized using HCR with DAPI stain in the mounting reagent. Scale bar represents 500um.

**Fig S5.**
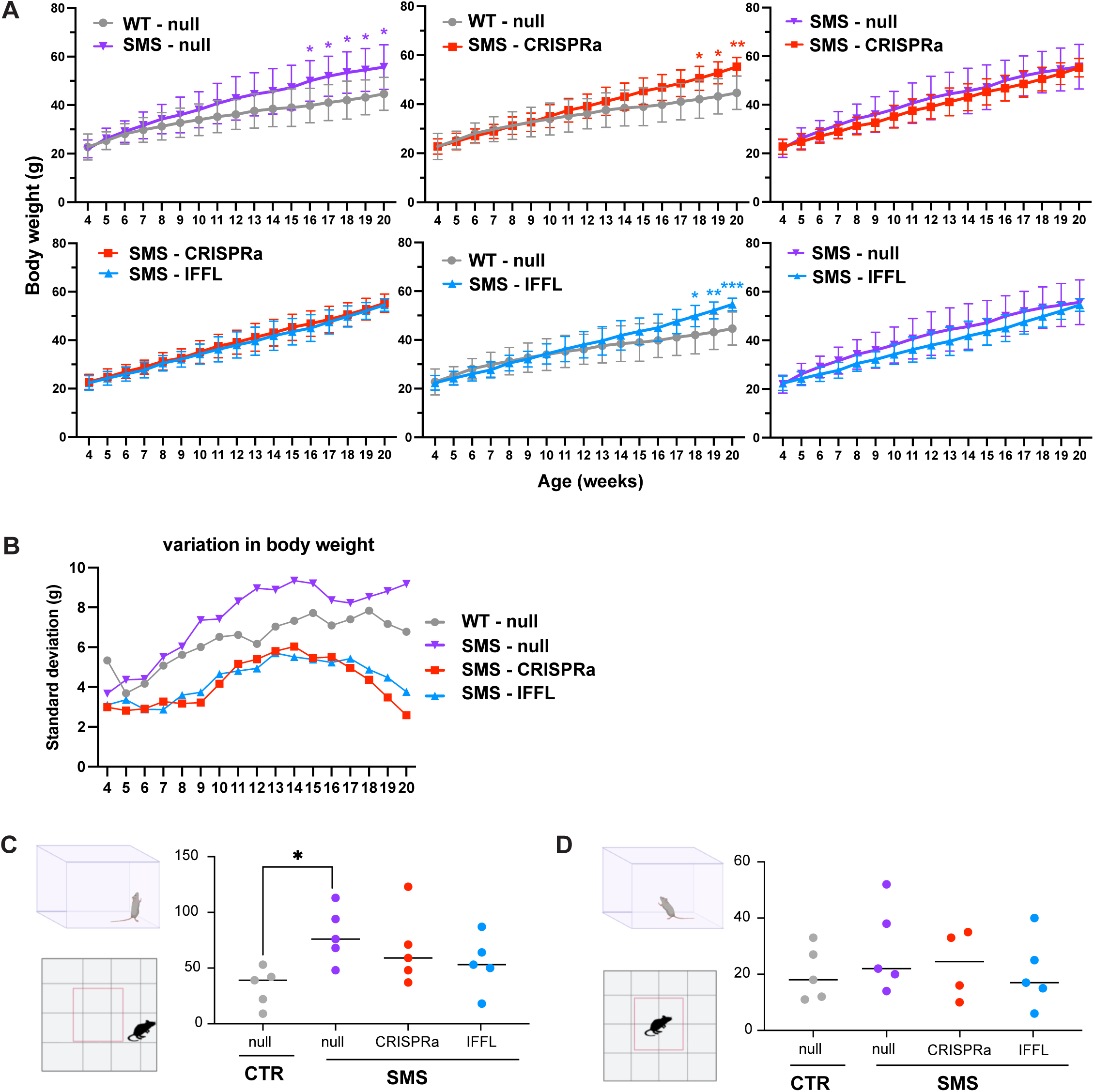
**A** Body weight comparisons of male mice across treatment groups ages 4-20 weeks. Two-way ANOVA followed by a Tukey’s multiple comparisons test was used to calculate p-value. **B** Standard deviation of body weight for male mice in each group ages 4-20 weeks. Levene’s test for equality of variances was used to calculate p-value. **C** Total rearing number in the periphery of the open field for male mice in each group. Ordinary one-way ANOVA followed by Tukey’s multiple comparisons test was used to calculate p-value. **D** Total rearing number in the center of the open field for male mice in each group. Ordinary one-way ANOVA followed by Tukey’s multiple comparisons test was used to calculate p-value (all comparisons were found to be not significant).

**Fig S6.**
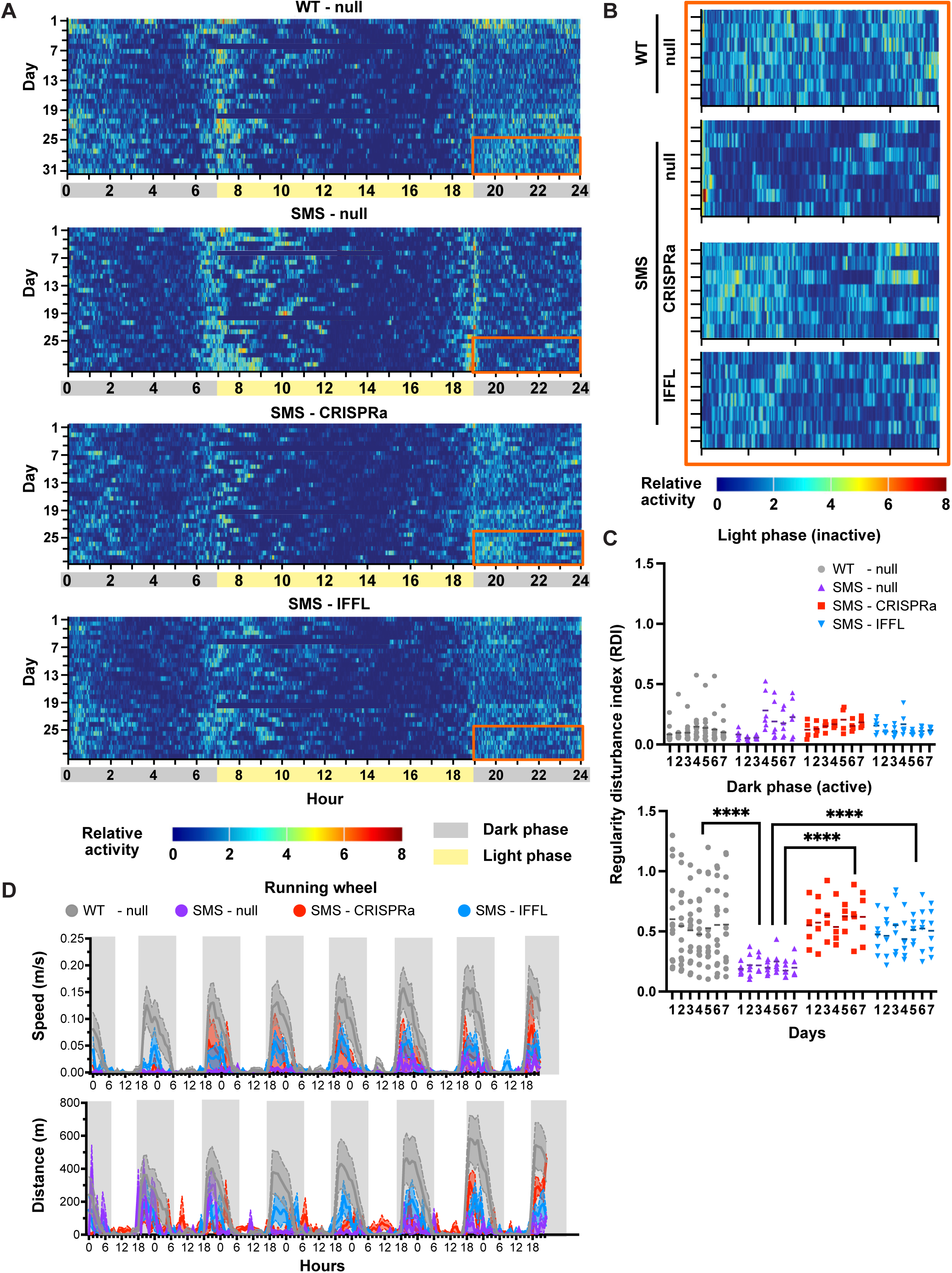
**A** Heat maps of average relative activity over four weeks for male mice in each group (WT-null, SMS-null, SMS-CRISPRa, SMS-IFFL). Representative dark phase activity during the last week when the mice are most acclimated is boxed in orange (see Fig6C). **B** Heat maps of representative dark phase activity of male mice in each group during the last week of DVC data collection (boxed area in Fig 6B). **C** Dot plot of light phase and dark phase regularity disturbance index of male mice in each group. Two-way ANOVA followed by Tukey’s multiple comparisons test was used to calculate p-value. **D** Mean and SEM of running wheel speed and distance of male mice in each group.

## Methods

### Construction of plasmids

Plasmids were generated using standard molecular cloning practices, including the following methods: In-Fusion, ligation, PCR, and annealing. All transgene sequences were ordered from Twist Biosciences or PCR amplified from plasmids (Addgene) using primers ordered from IDT and Thermo Fisher Scientific. Backbone plasmids were restricted using restriction enzymes from Fisher or NEB and purified. Then transgenes and backbones were assembled via in-fusion (Takara) or ligation (NEB), transformed via the heat-shock protocol into competent *E. coli* cells (Turbo-NEB), and plated on carbenicillin containing agar plates. Plasmids were extracted using Qiagen miniprep kits and verified by either Sanger sequencing or whole-plasmid sequencing (Pladmidsaurus). Plasmid sources for dCas9 and TRE-targeting gRNA were generous gifts from Prof. Stanley Qi at Stanford.

### Tissue culture

All cells were cultured in a humidity-controlled incubator under standard culture conditions (37°C with 5% CO2).

#### HEK293, COLO320, N2A, and CHO cells

HEK293 cells (catalog no. CRL-1573), and N2A cells (VWR, catalog no. MSPP-CCL131) were purchased from ATCC while COLO320 cells were a gift from the Howard Chang Lab and cultured using Dulbecco’s modified Eagle’s medium (Fisher, 501015428) supplemented with 10% fetal bovine serum (FBS) (Fisher Scientific catalog no. FB12999102), 1 mM sodium pyruvate (EMD Millipore catalog no. TMS-005-C), 1% penicillin-streptomycin (Genesee catalog no. 25-512), and non-essential amino acids (Genesee catalog no. 25-536). CHO cells were a gift from the Longzhi Tan Lab and cultured using F-12K nutrient mixture (Fisher, 21127022) supplemented with 10% FBS and 1% penicillin-streptomycin. The GFP-integrated HEK, COLO, and CHO cell lines were cultured in similar conditions with the addition of Hygromycin.

#### Lymphoblastoid cell lines (LCLs)

LCLs were purchased from the Coriell Institute (catalog no. GM23786) and cultured according to their protocol using RPMI medium supplemented with 10% FBS and 1% penicillin-streptomycin. They were grown in vessels for suspension cells with a breathable film cover. Cells were split once every three days and cultured at a 1:5 ratio in fresh media.

#### Mouse cortical neurons

Primary mouse neurons were purchased from Thermo Fisher Scientific (catalog no. A15585), thawed, and cultured according to recommended guidelines. Medium was prepared as follows: Complete Neurobasal™ Medium supplemented with GlutaMAX™ -I Supplement (Cat. no. 35050) and B-27™ Supplement (Cat. no. 17504) to final concentrations of 0.5mM and 2%, respectively. Plates were prepared as follows: 24-well plates were incubated with Poly-D-Lysine in PBS at 4.5 µg/cm2 for 2 hours in a biological hood, followed by 3x washes with water. Plates were then left open to dry for 1-2 hours and kept at 4C until use. Cells were thawed and cultured as follows: cells went through rapid thawing (< 2 minutes) of the frozen vial in a 37°C water bath, then transferred to the cell culture hood. Pipette tip was rinsed with complete medium and very gently transferred the cells to the pre-rinsed 50-mL tube 3 mL of complete medium (pre-warmed to 37℃) was added to the cells in the 50-mL tube extremely slowly at the rate of one drop per second. The suspension was then mixed and seeded in a pre-coated glass bottom 24-well plate (ibidi, 82426) with 1ml complete medium pre-warmed to 37°C at 200k cells per well. 50% of the media was changed 24 hours post-seeding and repeated every 2-3 days.

### Transient DNA transfections

Cells were cultured in 24-well tissue culture-treated plates under standard culture conditions. When cells were 60–90% confluent, they were transiently transfected with plasmids via the jetOPTIMUS DNA transfection reagent (Polyplus catalog no. 117-15), as per manufacturer’s instructions using 0.4 µl of reagent per 50 µl of jetOPTIMUS buffer for 500 ng total DNA transfections in the 24-well plate. To create GFP integrated cell lines, we transfected HEK293/CHO/COLO cells with NK90 piggybac plasmid (pTRE3-GFP) with a matching transposase, let them grow for 72 hours, and then added Hygromycin for selection. After 7 days, surviving cells were frozen and used in all relevant experiments.

### mRNA synthesis and transfection

The DNA templates for in vitro transcription contained dCas9 and TEV coding sequences flanked by optimal 5’ and 3’ UTRs as well as a T7 promoter and PolyA tail (NK409, NK411). The plasmid template with optimal UTRs was a gift from Prof. Michael Elowitz. In vitro transcribed mRNA was produced and purified by GeneScript Inc. based on the DNA template. gRNAs were produced based on Precision gRNA Synthesis Kit (Thermo Fisher Scientific, A29377). The purified mRNAs and gRNA were then transfected to HEK293-pTRE3G-EGFP at 90% confluency via the Mirus TransIT®-mRNA Transfection Kit following the manufacturer’s instructions. Briefly, 3 μl of TransIT-mRNA reagent as well as mRNA boost were mixed with mRNA and gRNA per well in 24 well plate. Expression of fluorescent proteins on the mRNA was assayed via flow cytometry 48 hours post transfection.

### Flow cytometry and data analysis

Cells were harvested approximately 48 h post transfection by trypsinization and resuspended in flow buffer (HBSS + 2.5 mg ml−1 bovine serum albumin). Cells were analyzed by flow cytometry (Biorad ZE5 Cell Analyzer), and data was processed with Matlab/FlowJo. Cells were gated for live and single cells based on FSC/SSC and FSC-A/FSC-W plots. They were then sorted according to the transfection marker signal and their output fluorescence signal averaged over a running window of 1%-5% of total count. Running averages of output fluorescence were then plotted on as a function of transfection marker signal. For HCR experiments, the fluorescent signal detected was altered to AlexaFluor-488, AlexaFluor-546, and AlexaFluor-647, for detection of GAPDH, BFP and Rai1, respectively.

### RNA extractions and quantitative PCR

Cells were grown as detailed above and were spun down at 500g for 5 minutes. RNA was then extracted using RNAasy mini kit (Qiagen), RNase-Free DNase Set (Qiagen), and QIAshredder (Qiagen). After extraction RNA concentration was measured via nanodrop. 3000 ng of purified RNA was then reverse transcribed using SuperScript™ IV First-Strand Synthesis System (Invitrogen). cDNA were then diluted and added to a qPCR compatible 96-well plate (Fisher, 4346907) together with qPCR master mix: SENSIMIX SYBR HI-ROX (VWR, 490017-866 / 868) and primers targeting human or mouse Rai1 gene. qPCR was carried out on a QuantStudio3 (Applied Biosystems) using SYBR-Green. RNA estimation was calculated based on the calibration curve of purified plasmid and normalized by the *Ct* threshold. The following primer pair sequences for mouse Rai1, human Rai1 and normalizing gene (β-actin) were used: Rai1-mouse-F, GCTCGTGACAGCTGGTACA; Rai1-mouse-R, AGGAACACTGGGTCCATGAG; Gapdh-Mouse-F, CATGGCCTTCCGTGTTCCTA; Gapdh-Mouse-R, CCTGCTTCACCACCTTCTTGA; RAI1-human-F, CCCAGGAGCACTGGGTGCATGA; RAI1-human-r, GCAGCTGGAACACATCATGTCCACG; β-actin-human-F, CGTCCACCGCAAATGCTT; β-actin-human-R, GTTTTCTGCGCAAGTTAGGTTTTGT.

### dCas9 *RAI1* gRNA screening

The sgRNA oligonucleotides targeting human RAI1 promoter regions were designed using the Benchling gRNA Design Tool.^57^ The sadCas9-2 × VP64 vector (Addgene #135338)^58^ was used as a backbone vector. It carries mutations in the endonuclease catalytic residues (D10A, N580A) of a FLAG-tagged saCas9, which was fused on both N and C termini with transcriptional activators VP64 (four copies of VP16). sgRNAs were then cloned into the backbone using the ligation cloning method previously described. After validating the sequences of constructs, they were transfected into 293A cells using polyethylenimine, and cells were harvested in TRIzol reagent 72 hours after transfection. Total RNA was extracted using phenol-chloroform extraction method, reverse-transcribed using the SuperScript III First-Strand Synthesis System (Thermo Fisher), and qPCR reactions were conducted using SsoAdvanced Universal SYBR Green Supermix (Bio-Rad) in StepOnePlus real-time PCR system (Applied Biosystems) with GAPDH as a housekeeping control.

The primers used for quantitative PCR were as follows: RAI1-human-F, CCTCAGCATTCCCAGTCCTTC; RAI1-human-R, CTGTGCAACTCTTATAGGAGTGG; GAPDH-human-F, AAGGTGAAGGTCGGAGTCAA; GAPDH-human-R, AATGAAGGGGTCATTGATGG.

### CasRx repression work

#### Determining the minimal collateral activity regime

To facilitate screening of cleavable CasRx designs, we use a synthetic system where we use the fluorescent protein mCherry as the output. Since it has been reported that CasRx can cleave non-target RNA, we first determine the regime in which this collateral activity is minimal. Since collateral activity for CasRx depends on target mRNA level, we titrate mCherry expression level (the target mRNA) by changing the promoter of the mCherry plasmid (CMVTO or SFFV) and the amount of plasmid transfected. We also measure GFP fluorescence which is expressed on the same transcript as CasRx (EF1a-GFP-T2A-CasRx) to determine the amount of off-target effect. Transfecting 50 ng of SFFV-mCherry plasmid resulted in the greatest fold change in mean mCherry fluorescence between control gRNA and mCherry-targeting gRNA and minimal difference between mean GFP fluorescence.

#### Repression circuit titration

Lastly, we used teD-cleav_CasRx7 in a whole circuit titration. We transfect 50 ng of SFFV-mCherry and varying amounts of the teD-cleav_CasRx7 circuit. We use BFP as a transfection marker and average the mCherry fluorescence for cells that are highly transfected (BFP fluorescence > 10^8^ A.U.) to produce Fig. 2F. Note that in the running average plots (Fig. S2F), the output mCherry fluorescence increases as dosage increases (BFP transfection marker) which is expected. This is because the mCherry gene is not integrated into the cell’s genome and is instead transcribed from a transfected plasmid. Thus, the relationship between mCherry fluorescence and dosage is not the same as in Fig. 2E which is derived from the simulation which assumes that the target gRNA is produced from an integrated gene.

### AAV production and transductions

AAVs were ordered and manufactured with the help of the Stanford Gene Vector and Virus Core (GVVC). Plasmids were amplified and extracted via the Qiagen plasmid midiprep kit (Qiagen), their concentration measured and sent to GVVC for production. The three AAVs for the dual AAV system were measured for their ITR: NK355: 1.12e13 vg/mL, NK404: 2.57e13 vg/mL, NK405: 2.05e13 vg/mL. For a 24-well of primary neurons, 4ul of total virus was with NK355 always in access, since dCas is the limiting factor and has a lower ITR. The single AAVs were measured for their ITR as well: NK404: 8.22e13 vg/mL and NK404: 7.49e13 vg/mL. Cells were monitored via the EVOS M7000 Cell Imaging System every day and were fixed 2.5 weeks after AAV transduction once a BFP signal was detected (if applicable). s

### Lentivirus production and transductions

LentiX cells were seeded in 6-well tissue culture plates one day prior to transfections. When cells were 80%-90% confluent, they were transiently transfected with plasmid constructs (600 ng PAX2, pMD2g, and 1,100 ng transgene plasmid) as detailed earlier. Cells were incubated for 24 h under standard culture conditions and were supplemented with 3ml of complete DMEM media to each well. Lentivirus was concentrated 24 hrs afterward using viral precipitation: for each lentiviral prep, media was filtered using a syringe and 0.45 µm filter into 15 mL conical tubes. 5x Lentivirus Precipitation Solution (Alstem 480 catalog no. VC100) was mixed with each prep and incubated at 4 °C for 48-72 hours. Viruses were then spun down at 1500xg for 30 min at 4 °C. Supernatant was aspirated, and virus was resuspended using 200 µL complete media (either DMEM for HEK293 and N2A, or RPMI for LCLs). Virus was then added to cells according to the following protocols, and leftover was frozen at -80°C for further use.

#### HEK293 and N2A cells

Virus was added dropwise onto 24-well tissue culture plates containing HEK293 or N2A cells seeded at 200k cells/well. Media was changed 24 hours post transduction, and cells were grown for an additional 5 days, at which time a positive BFP signal was detected, and then fixed and stained according to the following protocols.

#### Lymphoblastoid cell lines (LCLs)

Patient LCLs were split a day prior to transduction at a ⅓ ratio for optimal growth. On the day of transfection, cells were counted and 50k cells were transferred to multiple wells of a 96-well plate. Cells were then spun down at 500xg for 5 min and the media was gently taken out. Virus solutions were then supplemented by Polybrene (Merc) to a final concentration of 10ug/ml, and the mixed solutions were gently added to the cells without resuspension. Cells were then placed in the incubator overnight, and the virus solution removed by centrifugation (500xg for 5min). Cells were then merged and expanded, transferring from 96-well plates to a 48-well plate and finally a 24-well plate. Once cells reached confluency in 24-well plates, they were fixed and stained according to the following protocols.

#### Cell fixation and Hybridization Chain Reaction (HCR)

DNA probes for GAPDH, RAI1 and BFP were designed and ordered from Molecular Instruments based on the full-length genes with maximal available probe set size of ∼7500bp (Fig 3C, 3E, S3F). Gating was done based on positive probe signals, FSC-SSC for live cells, high GAPDH for active cells, and positive BFP mRNA signal for HCR-flow. All protocols were conducted according to Molecular Instruments HCR protocols. Briefly:

#### Cells in suspension: HEK293, N2A, B-LCL cells

**Day 1:** Growth media was aspirated from culture plates and cells were washed with DPBS. Cells were then trypsinized and quenched by the addition of growth media. These steps were omitted for suspension cells (B-LCLs). Cells were then transferred to 15 mL conical tubes and centrifuged for 5 min at 180xg. Cells were then resuspended in 4% formaldehyde to reach approximately 1 million cells/mL. Cells were fixed for 1 hr at room temperature, and then centrifuged and washed with PBST 4 times. Cells were then re-suspended in cold 70% ethanol overnight.

**Day 2:** Cells were centrifuged for 5 min and washed twice with 500 µL of PBST. PBST was then removed, and the pellets were resuspended with 400 µL of probe hybridization buffer and pre-hybridized for 30 min at 37°C. In the meantime, probe solution was prepared by adding a fixed amount of each probe set to 100 µL of probe hybridization buffer pre-heated to 37°C. Amounts used: 4 pmol for GAPDH, 16 pmol for RAI1, and 4 pmol for BFP. This solution was added to cells, and then they were incubated overnight (>12 h) at 37°C.

**Day 3:** Probe solution was removed, and the cells resuspended with 500 µL of probe wash buffer, incubated for 10 min at 37°C, and then centrifuged for 5 min to remove the probe wash buffer. This step was repeated three additional times. Next, the pellet was resuspended with 500 µL of 5× SSCT, incubated for 5 min at room temperature, then centrifuged and re-suspended with 150 µL of amplification buffer and let sit for 30 minutes at room temperature. Then the amplification hairpins were separately prepared: 15 pmol of hairpin h1 and 15 pmol of hairpin h2 for each primary hairpin (6 total) by heating to 95°C for 90 seconds and then cooled to room temperature in a dark drawer for 30 min. The hairpins were then added to the sample and incubated overnight (>12 h) in the dark at room temperature.

**Day 4:** Cells were then centrifuged for 5 min and hairpin solution removed. The cells were then washed with 5× SSCT six times. Finally, cells were resuspended with flow buffer, filtered, and analyzed via flow cytometry as detailed previously.

#### Cell on a slide: mouse cortical neurons

**Day 1:** Growth media was aspirated from culture plates and cells were washed with PBST twice. 4% formaldehyde was then added to the cells and fixed for 1.5 hr at room temperature. Next, cells were washed 2x times with PBST and incubated 10min with 0.3% Triton-X-100. Samples were then washed with 2 × 300 µL of 2× SSC and pre-hybridized in 300 µL of probe hybridization buffer for 30 min at 37°C. Probe solution was then prepared similarly to the previous protocol, except amounts of probes were increased to: 8 pmol for GAPDH, 32 pmol for RAI1, and 10 pmol for BFP. They were then added to probe hybridization buffer, placed on the cells, and left overnight (>12 hrs).

**Day 2:** Excess probes were removed by washing 4 × 5 min with 300 µL of probe wash buffer at 37°C. Then, samples were washed with 2 × 5 min with 5× SSCT at room temperature, and then pre-amplified in 300 µL of amplification buffer for 30 min at room temperature. Hairpin amplification solution was prepared similarly to the previous protocol, and then added to the cells. Cells were then left at room temperature for 2 hours, and then washed 5 × 5 min with 300 µL of 5× SSCT at room temperature. Finally, 5× SSCT was aspirated and 300ul of ProLong® Gold Antifade Reagent with DAPI was added (Cell Signaling). Samples were then stored at 4°C and protected from light prior to imaging. Imaging took place over multiple days starting at one day post storage.

### Brain slice fixation and HCR

#### Fixation

Mounted brain slices removed from -80°C and immersed in ice-cold 4% paraformaldehyde for 15 min at 4°C. The slides were immersed in 50%, 70%, and then 100% EtOH for 5 min at room temperature sequentially. Slides were immersed in another fresh 100% EtOH for 5 min at room temperature and then air dried for another 5 min at room temperature. 10 ug/mL proteinase K solution was introduced on top of the sample and incubated in a humidified chamber for 10 min at 37°C. Slides were washed by immersing in 1x PBS twice. Then a hydrophobic pen was used to draw a barrier around the samples.

#### HCR Gold RNA-FISH Protocol: Sample on a Slide

Brain slices were processed using the HCR Gold RNA-FISH protocol (for sample on a slide) immediately after the fixation protocol.

Probe hybridization: 200 µL of preheated (37°C) HCR™ HiFi Probe Hybridization Buffer was added on top of the sample and the sample was incubated at 37°C for 10 min inside a humidified chamber. Probe solution was prepared by adding HCR™ HiFi Probes (targeting *Gapdh*, *dCas9*, and *Rai1*) to HCR™ HiFi Probe Hybridization Buffer in a ratio of 4 µL probe to 100 µL buffer at 37°C. Pre-hybridization solution was removed and excess buffer drained on slide by blotting edges on a Kimwipe. Hybridization was performed by adding 50–100 µL of the probe solution on top of the sample, placing a coverslip on the sample, and incubating at 37°C overnight in the humidified chamber. The slide was then immersed in HCR™ HiFi Probe Wash Buffer at 37°C to float off coverslip. Excess probe was removed by immersing 4 × 15 min in HCR™ HiFi Probe Wash Buffer at 37°C. Slide was then dried by blotting edges on a Kimwipe.

Amplification: 200 µL of HCR™ Gold Amplifier Buffer was added on top of the sample and incubated at room temperature for 30 min for pre-amplification. Hairpins were prepared separately for each probe and snap cooled by heating at 95°C for 90 seconds and cooling to room temperature in a dark drawer for 30 min. Amplifier solution was prepared by adding snap cooled hairpins to the HCR™ Gold Amplifier Buffer. Then, the pre-amplification solution was removed and 50–100 µL of the amplifier solution added on top of the sample. A coverslip was placed on the sample and left to incubate at room temperature >3 h in the dark. Afterwards, slides were immersed in HCR™ Gold Amplifier Wash Buffer at room temperature to float off the cover slip. Excess amplifier was removed by immersing 4 × 15 min in HCR™ Gold Amplifier Wash Buffer at room temperature in the dark. Slides were dried using a Kimwipe. Antifade mounting reagent was added on top of the sample along with a coverslip. Samples were stored at room temperature for 24 hours then at 4°C and shielded from light prior to imaging.

### Microscopy experiments and analysis

#### Primary mouse cortical neurons

All images were taken with the Inverted Zeiss LSM 780 multiphoton laser scanning confocal microscope. The following channels were used: AlexaFluor-488, AlexaFluor-546, AlexaFluor-647, and DAPI. Z-stack captures of 10-30 slices for each field of view were used to capture the entire volume of cells. Images for the dual AAV experiment were analyzed using FIJI/ImageJ: all Z-stack slices were combined using the MAX pixel intensity (z-project). GAPDH and DAPI channels remained untouched, while the Rai1 channel went through thresholding of 0.4% of intensity to MAX intensity for each well, and outlier pixels were removed (1 pixel size and 5 minimum intensity). Images were then converted into masked images (binary). Finally, the three channels were then combined, and the outline of each cell was manually drawn based on the combined GAPDH + DAPI signals. The percentage of positive RAI1 pixels in each cell was then automatically calculated and documented. Data was collected from two individual wells and combined. Images for the single AAV experiment were also analyzed using FIJI/ImageJ with a similar protocol. However, the Rai1 channel was threshold based off 0.4% of the MAX intensity of the no transduction well and applied to all conditions. Data across wells was not combined.

#### Brain slices

Tissue sections were imaged on a Zeiss LSM900 with Airyscan (four channels: DAPI, AF546 [dCas], AF488 [GAPDH], AF647 [RAI1]), and z-stacks were reduced to single planes by maximum-intensity projection. Nuclei were segmented from the DAPI channel using Cellpose (v4, GPU-accelerated), applied to overlapping ∼800×800 px tiles rather than the full field of view to avoid under-segmentation of touching nuclei caused by global intensity normalization on larger images. Segmentation used a flow threshold of 0.6, and candidate nuclei smaller than 3,000 px² were discarded as segmentation artifacts. Whole-cell boundaries were assigned by marker-controlled watershed, seeded from nuclei and run over a grayscale-closed (morphological closing, disk radius 20) GAPDH channel to prevent local dips in GAPDH intensity from generating spurious watershed boundaries that fragment single cells. Cells whose resulting whole-cell mask did not expand beyond 8,000 px² (i.e., failed to grow appreciably past the nucleus) were excluded as likely segmentation failures.

Segmented cells were retained for quantification only if they met all of the following criteria: mean GAPDH intensity ≥1,000 (GAPDH-active, indicating a genuine cell rather than background); nucleus morphology and size consistent with a single, non-overlapping nucleus (area and solidity within a robust z-score-based threshold of the population median); and, for CRISPRa and IFFL constructs, mean dCas intensity ≥6,000 as a marker of highly transduced cells and effective construct delivery. Saturated, non-biological artifacts in the target-transcript channel (large, fully saturated, round blobs consistent with tissue debris) were identified by intensity- and size-based filtering (≥99.5th percentile intensity, ≥35 px equivalent diameter) and excluded from target-channel quantification only. Mean target-transcript intensity was computed per cell within the final whole-cell mask. Image analysis code was developed with assistance from Claude Code (Anthropic).

### Analysis of predicted HI/TS genes

Since copy number variations are rare mutations, there is insufficient clinical data to understand dosage sensitivity for almost all genes. To address this problem, Collins et al. analyzed copy number variation data from 950,278 individuals and used machine learning to predict haploinsufficiency and triplosensitivity for all autosomal protein-coding genes within the human genome.^21^ In this paper, we use data from Table S7: Haploinsufficiency and triplosensitivity predictions for all autosomal protein-coding genes. First, we screen for genes which have both a haploinsufficiency score (pHaplo) >= 0.86 and a triplosensitivity score (pTriplo) >= 0.94, indicating that the genes are predicted to be both haploinsufficient and triplosensitive (as defined by the authors). This resulted in a total of 915 genes after screening. Next, we used the Proteins REST API which provides data from UniProt to determine the protein sequence for each gene.^59^ There were two genes with no return (AP000783.1, AC008443.1) which reduces the total number of dosage sensitive genes to 913. Lastly, we took the length of each protein sequence and multiplied by three to calculate the shortest possible gene length for each gene, which is an underestimate of the final sequence which would be necessary for proper expression that would need to fit within an AAV.

### Wild-type mouse husbandry and brain processing

8-week-old male C57BL/6 (0664) mice were purchased from The Jackson Laboratory. The animals were group-housed (four/five per cage) in an environmentally controlled room with a reverse 12-h light/dark cycle. Before surgery, mice were acclimatized to the housing facilities for at least a week. Mice were randomized into treatment groups based on cages. Mice in the same cage received the same treatment, except the last cage which included a mixture of treatment groups. Mice were anesthetized using 4% isoflurane (Isothesia, Henry Schein Animal Health, Dublin, OH) in a mixture of oxygen and compressed air. A toe pinch reflex confirmed full anesthetic induction. Immediately after induction, mice received a subcutaneous (s.c.) injection of 5 mg/kg Carprofen (Vet One Ostifen Injection). Mice were shaved on top of the skull and positioned on a digital stereotaxic frame (David Kopf Instruments), with isoflurane maintained at 1–2.5% throughout the surgery. Eye drops were applied to keep the cornea moisturized, and the skull surface was disinfected with Betadine and 70% ethanol solution. A median incision above the skull was topically anesthetized with 5 mg/mL lidocaine (Xylocaine MPF, Fresenius Kabi, USA). Respiration rate, skin color, and toe-pinch reflex were monitored every 15 min, while normothermia was maintained through a heating pad (WPI, ATC2000). Burr holes were drilled with size 007 bits from Fine Science Tools. Stereotaxic injection coordinates were AP = -0.55, ML=+/-0.3, DV-5.0 from ideal bregma. The equipment for the injection consisted of a Hamilton Neuros 700 Series Microliter Syringe with Removable Needle, a 33-gauge blunt Hamilton needle, and a microinjection pump (UMP3, WPI) connected to an intelligent touch controller (WPI). Mice were bilaterally infused with treatment at a rate of 0.1μL/min for a total of 0.5uL in each site. After the infusion, the needle was left in place for 2 min before being slowly withdrawn. The incision site was closed with 5-0 monofilament absorbable suture (421A).

After injection, mice were placed in recovery cages and kept warm with heating pads until ambulatory and awake before returning to their home cage. Carprofen was administered subcutaneously for 2 days following surgery day.

Four weeks after injection, mice were anesthetized and brains removed, embedded in OCT, frozen with isopentane and dry ice, and stored at -80°C. Coronal sections of the PVH regions were made at 15-20um thickness, mounted onto slides, and stored at -80°C.

### Mouse models and husbandry

All procedures were performed in accordance with the guidelines of the Canadian Council on Animal Care and the Montreal General Hospital Facility Animal Care Committee, with the appropriate approved protocols for animal use. Both male and female mice were used for the experiments. Mice were housed in groups on a 12-h light/12-h dark cycle with ad libitum access to food and water. The number of animals used in each experiment is provided in the text and figure legends. B6/CD1 F1 hybrid mice were used for all experiments. Rai1flox/flox mice (RRID:IMSR_JAX: 029103) were previously generated and characterized by our group.^26^ ActbCre/Cre mice were obtained from The Jackson Laboratory (RRID:IMSR_JAX: 033984).

#### Mouse genotyping

Genotyping for ActbCre mice was performed using primers 5′-CACCCTGTTACGTATAGCCG-3′ and 5′-GAGTCATCCTTAGCGCCGTA-3′ for a 300-bp Cre band, and primers 5′-CCAATCTGCTCACACAGGATAGAGAGGGCAGG-3′ and 5′-CCTTGAGGCTGTCCAAGTGATTCAGGCCATCG-3′ for a 500-bp internal control band.

Rai1flox mice were identified using the following primers: 5′-CAGAGTCCAGATGGCACTACAGGGG-3′, 5′-GTGAGCTCCCGCTGAAATGGACAGT-3′, and 5′-GGAGGTCTGCGCTTCAGGGCTTAAT-3′. With these primers, the wild-type Rai1 allele produces a 396-bp band and the Rai1flox allele produces a 497-bp band.

#### Neonatal intracerebroventricular AAV injection

Bilateral intracerebroventricular (ICV) injections were performed in neonatal (P0) mice as previously described^49,60^, relying on the ability of adeno-associated virus (AAV) to move freely from the cerebral ventricles into the brain parenchyma while the ependymal lining is still immature during the first 24 hr after birth. Litters were checked for birth at least twice daily, and pups were injected within 24 hr of delivery. Pups were anesthetized by hypothermia on a chilled metal plate for 2–3 min; anesthesia was confirmed by loss of the pedal withdrawal reflex and absence of spontaneous movement.

For the dual-AAV IFFL group, AAV encoding dCas9–zipper-cTEV and AAV encoding the Rai1 gRNA–zipper-nTEV cassette were combined at a 1:1 ratio in ice-cold PBS and co-injected so that proteolytic reconstitution of active TEV, and hence release of the CRISPRa activation domain, occurred only in cells transduced by both vectors. SMS-CRISPRa animals received the CRISPRa-only vector alone, and WT-null AAV and SMS-null AAV groups received a non-targeting control vector, all at matched total titer.

Injections were delivered bilaterally into the lateral ventricles using a 32G needle mounted on a microliter syringe, targeting a site approximately two-fifths of the distance from the lambda suture to each eye. The needle was advanced to a depth of ∼3 mm, and a slight decrease in resistance was used to confirm ventricular entry; targeting accuracy was validated in pilot pups by co-injecting 0.05% trypan blue and confirming dye spread throughout the ventricular system. A total of up to 2 µl of viral solution was injected per hemisphere, delivered slowly and without repuncturing a given injection site, before the contralateral ventricle was injected using the same procedure.

Following injection, pups were returned to a warming pad until normal coloration and movement resumed, then returned to the biological dam. Maternal retrieval and nursing were confirmed within 10 min.

#### Body weight measurement

The body weight of animals was measured weekly from week 4 (post-weaning) onward. After the 20th week, mice were transferred to digital ventilated cages (DVC) for home-cage activity monitoring.

#### Vertical rearing (open field) assay

To assess vertical rearing, mice were tested in an open-field arena at 8-10 weeks of age. On the day of testing, animals were transported to the testing room and habituated to the room for 1 hr prior to the start of the session. Each mouse was then placed in the center of a square open-field arena (45 × 45 × 40 cm [W × D × H]) and allowed to explore freely for 10 min while its behavior was video-recorded. Rearing events were classified as either periphery rearing, in which the forepaws contacted the arena wall, or center rearing, in which the mouse reared without wall support, and were quantified by an observer blind to genotype and treatment. The arena was cleaned with 70% ethanol between animals to eliminate residual olfactory cues.

#### DVC analytics

##### Home-Cage Activity Monitoring

Mice were individually housed in Digital Ventilated Cages (DVC®) throughout the recording period. Home-cage locomotor activity was recorded continuously over a 3–4-week period, in 1-min bins across the full 24-h cycle, using the DVC system (Tecniplast S.p.A., Buguggiate, Italy). Cages (GM500) were mounted on a standard individually ventilated cage rack instrumented with an external capacitance-sensing board positioned beneath each cage, such that no modification to or intrusion into the home-cage environment was required. Each board comprises 12 uniformly distributed, contactless electrodes whose electrical capacitance is sampled at sub-second intervals. Owing to the high water content of animal tissue, the presence and movement of mice near an electrode alters its measured capacitance; an electrode is registered as activated when the difference between consecutive capacitance measurements exceeds a fixed threshold (λ), selected to discriminate animal movement from noise-induced capacitance fluctuation, as previously described^31^. Data were processed and computed on the DVC Analytics platform (Tecniplast).

Electrode activation events were aggregated into 1-min bins to yield a continuous locomotor activity time series, hereafter referred to as activity, spanning the full 3–4-week recording period. For analyses of immobility, a lower activation threshold (λ = 1.25, applied to consecutive 1-s averaging windows) was used in order to exclude micro-movements and thereby distinguish complete immobility from slow or small-amplitude motion.^31^

##### Regularity Disruption Index

Irregularity of the activity pattern was quantified using the Regularity Disruption Index (RDI), a digital biomarker previously developed and validated in the SOD1G93A mouse model of amyotrophic lateral sclerosis^51^. RDI is derived from the sample entropy of the minute-based activity time series^61^, computed with embedding dimension m = 2 and tolerance r = 0.2. Prior to computation, activity values below a fixed threshold (λ = 0.005) were set to zero and a fourth-order Butterworth band-pass filter (normalized cut-off frequencies f_low = 1/2000, f_high = 1/300) was applied independently to each 24-h period; band-pass filtration was included because it yields more stable estimates than unfiltered data. RDI was computed over the 12 h of the lights-on phase and, separately, over the 12 h of the lights-off phase (N = 720 min in each case) for each day of the 3–4-week recording period. In addition, a central moving RDI was computed with a 60-min sliding window (N = 60), such that for each minute t the RDI was calculated over the 1-h interval centred on t, allowing the index to be visualized across the 24-h cycle.

RDI quantifies the irregularity of a time series and is independent of the absolute amount of activity it contains. A period in which successive minutes exhibit comparable activity levels yields an RDI approaching zero, whereas a period in which minute-to-minute activity varies markedly yields a high RDI. The index is scale-invariant: multiplying activity levels by a constant does not alter RDI.

##### Running Wheel Activity

Voluntary wheel-running was recorded using running wheels (DVC® GYM500, Tecniplast S.p.A.) fitted to each individually housed home cage. Wheel rotations were detected via an integrated magnetic sensor. Running distance was calculated by multiplying the number of complete wheel rotations by the wheel’s known perimeter (34.54 cm), and running speed was derived from the rotation rate over the same interval. Both measures were aggregated into 1-h bins over 1 week continuous recording periods. Bout structure, wheel occupancy time, and turn laterality were not analyzed in this study.

#### Mathematical modeling for activation circuit simulations

To predict the behavior of the activation feedforward circuit, we construct an ordinary differential equation model assuming Michaelis-Menten kinetics, steady-state, and quasi-equilibrium. We incorporate interactions between dCas9 and the DNA, mRNA and protein production, first-order degradation of mRNA and proteins, and dCas9 cleavage by TEV.

We define the probability of active dCas9 binding to the target DNA (e.g. the *RAI1* gene) as:

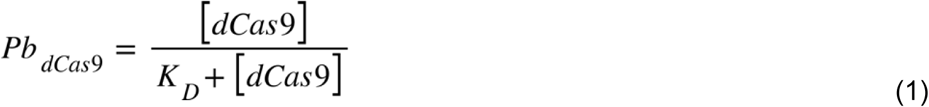

where *K_D_* is the dCas9 dissociation constant and [*dCas9*] is the concentration of the active dCas9 protein. When we assume that cleaved dCas9 species competes with active dCas9 species to bind to the DNA, we add another term in the denominator to account for this competitive inhibition:

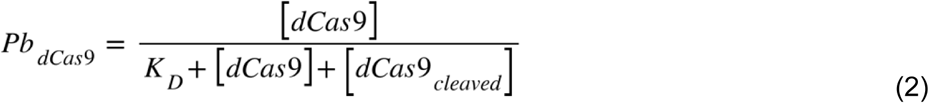

Where [*dCas9_cleaved_*] is the concentration of the cleaved dCas9 species.

Therefore, the change in corresponding target mRNA concentration is described by:

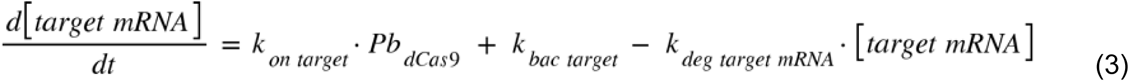

where *k_bac target_* is the background transcription rate of the target gene, *k_on target_* is the transcription rate when dCas9 is bound, and *k_deg target mRNA_* is the degradation rate of the target mRNA. The change in corresponding target protein concentration is described by:

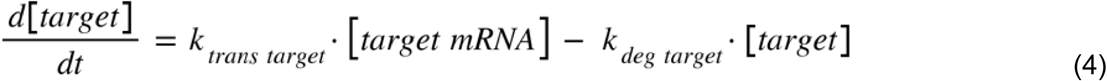

where *k_trans target_* is the translation rate of the target protein and *k_deg target_* is the degradation rate of the target protein. The steady-state solutions for equation (3) and (4) are:

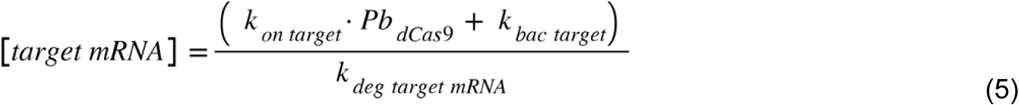

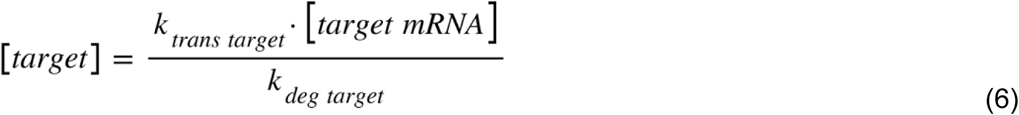

The dCas9 and TEV components are delivered to the cell such as in plasmid transfection and are therefore a function of dosage (DNA delivered). The TEV mRNA transcription rate can be represented by:

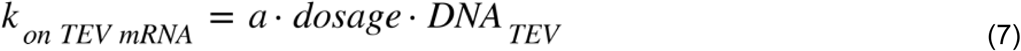

where *a* is a constant representing the TEV transcription rate in units of concentration/(time*plasmid amount) (e.g. nM/h-ng plasmid), *dosage* is the amount of plasmid that is delivered to the cell, and *DNA_TEV_* is the fraction of plasmid that encodes for TEV. The change in TEV mRNA concentration can therefore be described by:

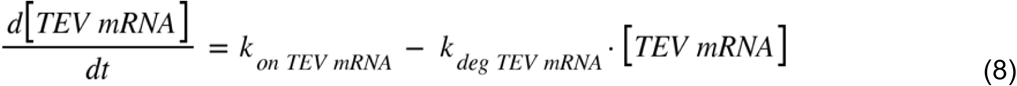

Similarly, the dCas9 mRNA transcription rate is represented by:

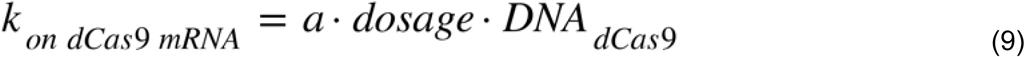

and the change in dCas9 mRNA concentration is described by:

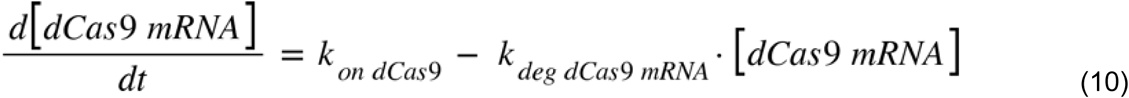

The change in TEV protein concentration is described by:

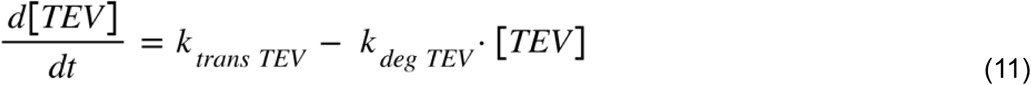

Lastly, the change in active dCas9 protein concentration is described by:

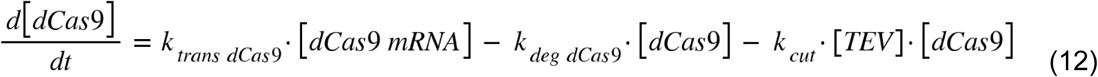

where *k_cut_*is the TEV *k_cat_/K_M_*, while the change in cleaved dCas9 protein concentration is described by:

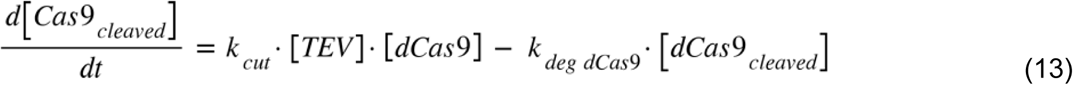

The steady-state solutions for equations (8), (10), (11), (12), and (13) are respectively:

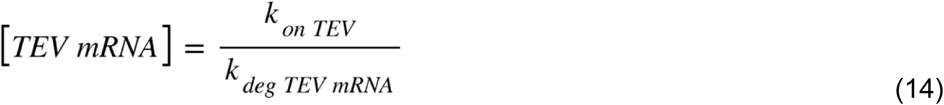

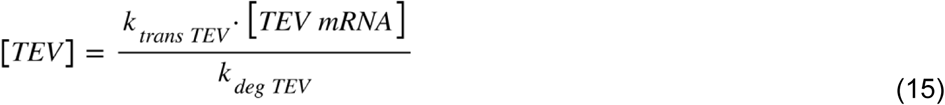

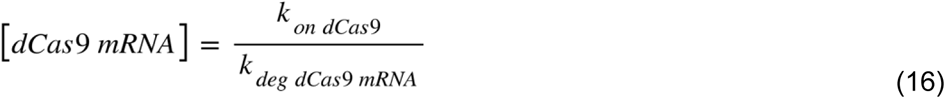

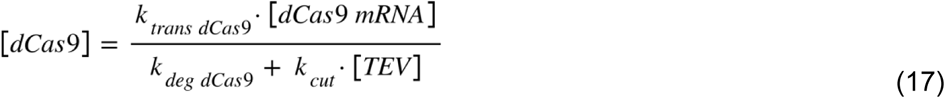

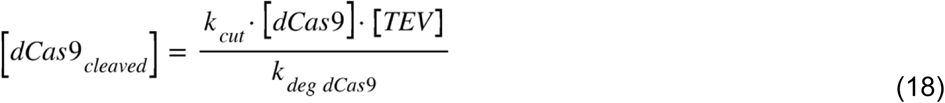

All numerical values, units, and references used for the parameters are found in Table S1.

### Gating strategy for activation circuit titration

We observe a non-monotonic trend at high transfection levels for the running means of the activation circuit titration (Fig S1B). Measuring the average GFP fluorescence of all transfected cells masks this non-monotonic effect (Fig S1A). Since this non-monotonic effect biases the average GFP fluorescence of all transfected cells, we present the activation circuit titration in **Fig 1F** as the average of transfected cells with an mCherry fluorescence below 10^7^.

### Engineering a TEV cleavable Cas13

#### Mathematical modeling for repression circuit simulations

To predict the behavior of the repression feedforward circuit, we construct an ordinary differential equation model assuming Michaelis-Menten kinetics, steady-state, and quasi-equilibrium. We incorporate interactions between Cas13 and the target mRNA, mRNA and protein production, first-order degradation of mRNA and proteins, and Cas13 cleavage by TEV. Unfortunately, there is little kinetic data for CasRx in contrast to TEV. Furthermore, much of the kinetic data available measures off-target mRNA cleavage rather than on-target mRNA cleavage. Therefore, we also assume that k_cat_/K_M_ for Cas13 is the same as TEV, though this is likely not the case. The change in corresponding target mRNA concentration is described by:

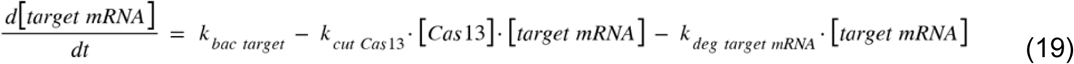

where *k_bac target_* is the background transcription rate of the target gene, *k_cut Cas13_* is the *k_cut_/K_M_* for Cas13, and *k_deg target mRNA_* is the degradation rate of the target mRNA. The change in corresponding target protein concentration is described by:

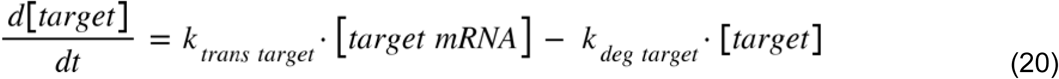

where *k_trans target_* is the translation rate of the target protein and *k_deg target_* is the degradation rate of the target protein. The steady-state solutions for equation (19) and (20) are:

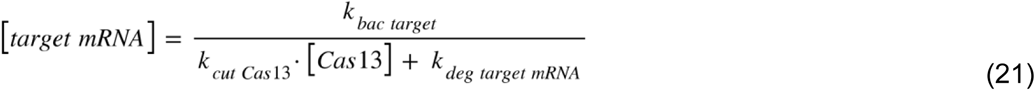

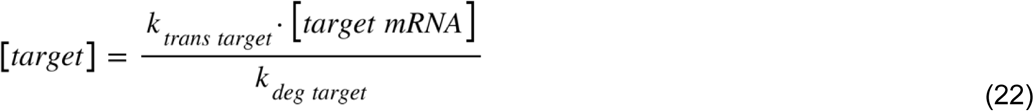

The Cas13 and TEV components are delivered to the cell such as in plasmid transfection and are therefore a function of dosage (DNA delivered). The TEV mRNA transcription rate can be represented by:

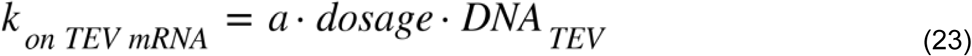

where *a* is a constant representing the TEV transcription rate in units of concentration/(time*plasmid amount) (e.g. nM/h-ng plasmid), *dosage* is the amount of plasmid that is delivered to the cell, and *DNA_TEV_* is the fraction of plasmid that encodes for TEV. The change in TEV mRNA concentration can therefore be described by:

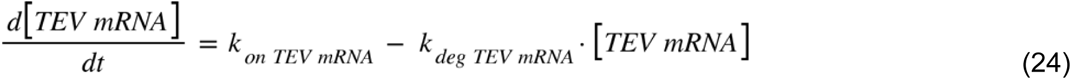

Similarly, the Cas13 mRNA transcription rate is represented by:

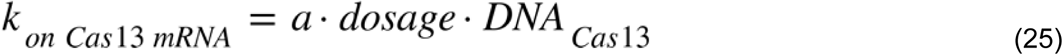

and the change in Cas13 mRNA concentration is described by:

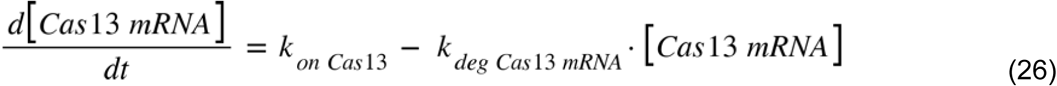

The change in TEV protein concentration is described by:

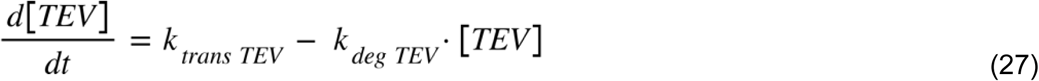

Lastly, the change in active Cas13 protein concentration is described by:

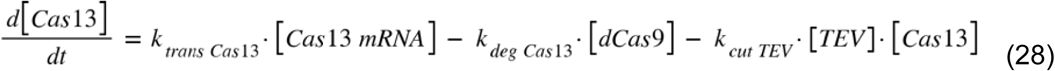

where *k_cut TEV_* is the TEV *k_cat_/K_M_*.

The steady-state solutions for equations (24), (26), (27), and (28) are respectively:

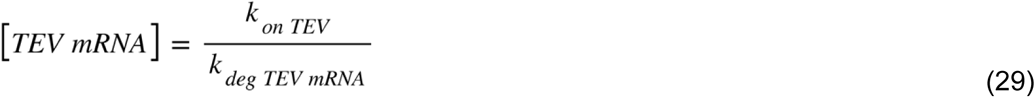

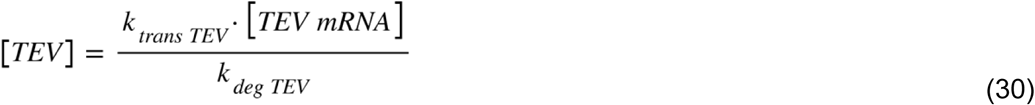

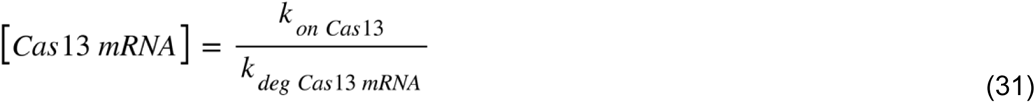

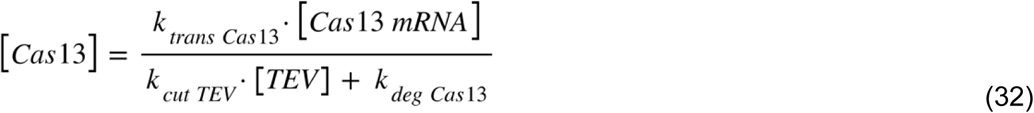

All numerical values, units, and references used for the parameters are found in Table S1.

### Calculating the plasmid transcription rate for the simulations

To estimate the value of the plasmid transcription eate *a*, as mentioned in equation (7), (9), (23), and (25), we use experimental data and data reported for GFP fluorescence in the supplementary information of a paper previously published by the Gao lab.^62^ We conducted an experiment where 10 ng of CMVTO-GFP was transfected for a single 24-well of HEK293 cells and measured an average GFP fluorescence of 1.3*10^6 A.U. From Fig. S2A and S2C from Kaseniit *et al*., we estimate that 300 copies of GFP mRNA corresponds to a measured fluorescence of 7*10^5 A.U. (we multiply the reported value by 100 due to differences in normalization between analysis software). 300 copies of mRNA is 0.5 nM of mRNA using Avogadro’s number and an estimated cell volume of 1 pL. Assuming a linear relationship between average GFP fluorescence and ng of DNA transfected, we would estimate that an average GFP fluorescence of 7*10^5^ would correspond to 5.38 ng of plasmid transfected. Since these measurements were taken at steady state, we calculate *a* using the following equation set equal to zero:

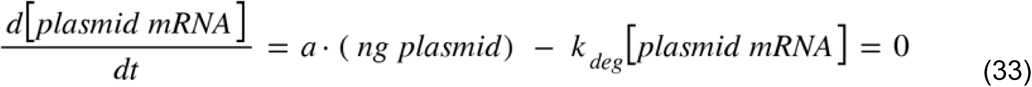

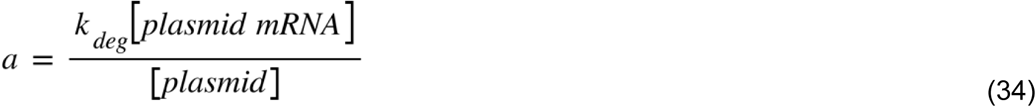

We estimate the ng plasmid per cell by dividing 5.38 ng by 0.24*10^6^, the number of cells within a confluent well of a 24-well plate,^63^ to determine (*ng plasmid*). We previously calculated [*plasmid mRNA*] to be 0.5 nM. Lastly, we use an estimate for the median mRNA degradation rate (Table S1). We then solve for *a* resulting in 1.71*10^3^ nM/(h-ng plasmid).

## Notes

https://data.mendeley.com/datasets/hxdzhsjvf7/1

### Competing Interest Statement

X.J.G. is a co-founder and serves on the scientific advisory board of Radar Tx. All other authors declare no conflict of interest.

### Summary of Updates

We have included experiments further characterizing the performance and behavior of our proteolytic IFFL in vitro and in vivo. We then deliver our circuit to a mouse model of Smith-Magenis syndrome and compare its performance to conventional CRISPRa.

